# Coordination of Pickpocket ion channel delivery and dendrite growth in Drosophila sensory neurons

**DOI:** 10.1101/2022.05.19.492359

**Authors:** Josephine W. Mitchell, Ipek Midillioglu, Ethan Schauer, Bei Wang, Chun Han, Jill Wildonger

## Abstract

Sensory neurons enable an organism to perceive external stimuli, which is essential for survival. The sensory capacity of a neuron depends on the elaboration of its dendritic arbor and the delivery of sensory ion channels to the dendritic membrane. However, it is not well understood how ion channels are trafficked to sensory dendrites and whether their delivery is coordinated with dendrite growth. We investigated the trafficking of the DEG/ENaC/ASIC ion channel Pickpocket (Ppk) in peripheral sensory neurons in fruit fly larvae. We used CRISPR-Cas9 genome engineering to tag endogenous Ppk1 and visualize it live, including monitoring Ppk1 membrane localization via a novel secreted split-GFP approach. Strikingly, Ppk1 is present throughout the membrane of actively growing dendrites, and Ppk1 density scales in proportion to the dendritic membrane, even when dynein-mediated transport to dendrites is disrupted. Our data suggest that Ppk1 is integral to the membrane of growing dendrites and implicate the recycling endosome GTPase Rab11 in the forward trafficking of Ppk1 to dendrites. Together, our results suggest that Ppk channel transport is coordinated with dendrite morphogenesis, thus ensuring proper ion channel levels and distribution in sensory dendrites.

**Author Summary:** Peripheral sensory neurons are essential for an organism to interact with its environment. Neurons are composed of signal-receiving dendrites and a signal-sending axon. Ion channels distributed throughout sensory dendrites transduce external stimuli into chemical signals, however the mechanisms that localize ion channels to sensory dendrites are not well understood. Both the composition of ion channels in the dendrites and the structure of a sensory neuron’s dendritic arbor are important for how it functions to sense external stimuli. Using live imaging and genomic engineering, we have discovered that the localization of a sensory ion channel, Pickpocket, in fruit fly sensory neurons is coordinated with growth of the dendritic arbor and that Pickpocket levels scale in proportion to dendrite length, even when transport to dendrites is disrupted. We also developed a novel genetically encoded approach to visualize the membrane expression of proteins in a living organism utilizing the split-GFP system. We found that both the secretory and endosomal networks mediate the forward trafficking of Pickpocket during neuronal morphogenesis, thus coordinating membrane growth with ion channel delivery. Our findings reveal that actively growing sensory dendrites are equipped with the ion channels needed for sensing external stimuli.

## Introduction

An organism’s interactions with its environment rely on its ability to sense external stimuli through sensory neurons. Ion channels distributed throughout the dendritic arbor of a sensory neuron rapidly transduce external stimuli into cellular signals. Both the morphology of a sensory neuron’s dendritic arbor and the localization of ion channels in the arbor are essential to the establishment of a neuron’s receptive field and sensory capacity. While the localization of ion channels to synapses in the central nervous system has been well studied [1], little is known regarding mechanisms that regulate the delivery of ion channels to the dendritic membrane of sensory neurons in the peripheral nervous system. It is also not known whether and how this trafficking may be coordinated with dendrite morphogenesis to establish the proper distribution of ion channels needed for sensing environmental stimuli.

To investigate the relationship between ion channel trafficking and dendrite growth, we used the *Drosophila melanogaster* class IV dendritic arborization (da) neurons as a model. The class IV da neurons function as polymodal nociceptors that detect multiple stimuli (thermal, mechanical, and light) and extend elaborately branched dendritic arbors that cover the larval body wall [2–5]. These neurons are an ideal model to study ion channel delivery in growing sensory dendrites for several reasons. First, during larval development, the class IV da neuron dendrites undergo expansive growth that can be easily visualized live in intact animals due to their superficial location just beneath the transparent larval cuticle and their relatively flat, two-dimensional morphology [6,7]. Second, the class IV da neurons have been a powerful *in vivo* model to identify mechanisms of dendrite morphogenesis, including players involved in membrane production and trafficking, the secretory and endosomal networks, molecular motor-based transport, and the cytoskeleton [8,9]. By manipulating known mechanisms of dendrite arbor growth, we can investigate how ion channel trafficking is coordinated with dendrite morphogenesis. Third, the general morphology and function of the class IV neurons is similar to peripheral sensory neurons and nociceptors in other organisms, including the mammalian C- and Aδ-fibers and the worm PVD and FLP neurons [10–12]. Thus, studying ion channel trafficking in the class IV da neurons may shed light on conserved mechanisms of ion channel localization in sensory dendrites.

During neuronal morphogenesis, the class IV da neurons express several dendritic ion channels that have been structurally and functionally characterized, including Pickpocket (Ppk); Transient Receptor Potential (TRP) channels, such as painless; and Piezo [3,13–16]. Whereas TRP and Piezo channels are comprised of large multi-pass membrane protein subunits, the Ppk ion channel subunits are relatively small, two-pass membrane proteins. Their modest size makes endogenous Ppk channels amenable to manipulation via CRISPR-Cas9 genome engineering. Moreover, the crystal structure of a conserved Ppk ortholog in chickens, ASIC1, has been solved, and this information is advantageous for the structure-guided manipulation of Ppk [17]. For these reasons we decided to focus on investigating the trafficking of Ppk in the class IV da neurons.

Ppk proteins belong to the large, structurally conserved family of Degenerin/Epithelial Na^+^ Channel/Acid Sensing Ion Channels (DEG/ENaC/ASICs) whose members in worms, flies, fish, and mammals carry out a variety of functions ranging from mechanosensation and learning and memory in the nervous system to salt homeostasis in epithelial cells in the kidney [18–20]. In the fly class IV da neurons, the Ppk channel is composed of two subunits, Pickpocket 1 (Ppk1) and Pickpocket 26 (Ppk26), which are mutually dependent on each other for membrane expression [16,21–23]. The localization of Ppk1 and Ppk26 has been characterized using antibodies and fluorescently tagged transgenes, and both subunits are broadly distributed throughout the developing dendrites of class IV da neurons. Interestingly, Ppk1 and Ppk26 are expressed from late-embryo to mid-larval stages, which coincides with a period of rapid dendrite growth [6,16,21,24–26]. The timing of Ppk1 and Ppk26 expression suggests that Ppk channel production and localization may be coordinated with dendrite growth.

To investigate the potential coordination of Ppk1 trafficking and dendrite growth, we used CRISPR-Cas9 genome engineering to tag endogenous Ppk1 and follow its localization in growing dendrites. We found that in developing neurons, Ppk1 is enriched throughout dendrites and is also present in axon terminals and at low levels in axons. Using a new split-GFP strategy to monitor the insertion of proteins into the cell membrane, we visualized Ppk1 membrane expression live in developing neurons, which revealed the robust, uniform membrane localization of Ppk1 in the somatodendritic compartment. We found that Ppk1 was present throughout growing dendrite branches and in actively growing dendrite tips, suggesting that Ppk1 is part of the nascent membrane that is adding to growing dendrites. In support of this model, we found that Ppk1 dendritic levels scale in proportion to the amount of dendritic membrane, even when transport to dendrites is disrupted by perturbing dynein. We also discovered that the recycling endosome GTPase Rab11 is involved in forward trafficking of Ppk1 to dendrites, which indicates a role for endosome-mediated trafficking in both dendrite growth and the delivery of a sensory ion channel. Together, our results suggest that Ppk channel delivery is coordinated with sensory dendrite morphogenesis, thus revealing a mechanism to establish proper ion channel levels and distribution throughout sensory dendrites.

## Results

### Ppk1 is enriched in dendrites and the dendritic membrane in developing sensory neurons

To visualize the localization of the Ppk ion channel in sensory neurons, we tagged endogenous Ppk1 with fluorescent proteins. To facilitate the manipulation of *ppk1*, we first replaced the ppk1 gene with an attP “docking site,” which enables the reliable and rapid knock-in of new *ppk1* alleles (Fig 1A). We then used this strain to knock-in *ppk1* tagged with one copy of superfolder GFP (sfGFP) at either the N- or C-terminus (Fig 1B and S1 Fig; since Ppk1 tagged with GFP at either terminus displayed similar localization, we used C-terminally tagged Ppk1 for most of our experiments). Excitingly, we observed fluorescent signal in neurons in live animals with just one copy of GFP attached to Ppk1 (Fig 1B). Consistent with previous reports, Ppk1::sfGFP was expressed in the class IV da neurons in the peripheral nervous system [2,16,25]. In the dorsal class IV da neuron called ddaC, Ppk1::sfGFP localized to both dendrites and axons, but its distribution to and within these compartments differed. Ppk1::sfGFP was enriched in dendrites, where it appeared to localize predominantly to the dendritic membrane and was present throughout the dendritic arbor (Fig 1B). This distribution matches the previously reported distribution of Ppk1 based on antibody staining and fluorescently tagged Ppk1 transgenes [21,23,27]. In contrast to dendrites, the Ppk1::sfGFP signal was dimmer in axons and did not align with the axonal membrane (Fig 1B). In the ventral nerve cord, where the ddaC axons terminate, Ppk1::sfGFP was present in axon terminals (Fig 1C). Altogether, our data indicate that Ppk1::sfGFP localizes predominantly to dendrites, but that it is also present at low levels in axons and in axon terminals.

**Fig 1.**
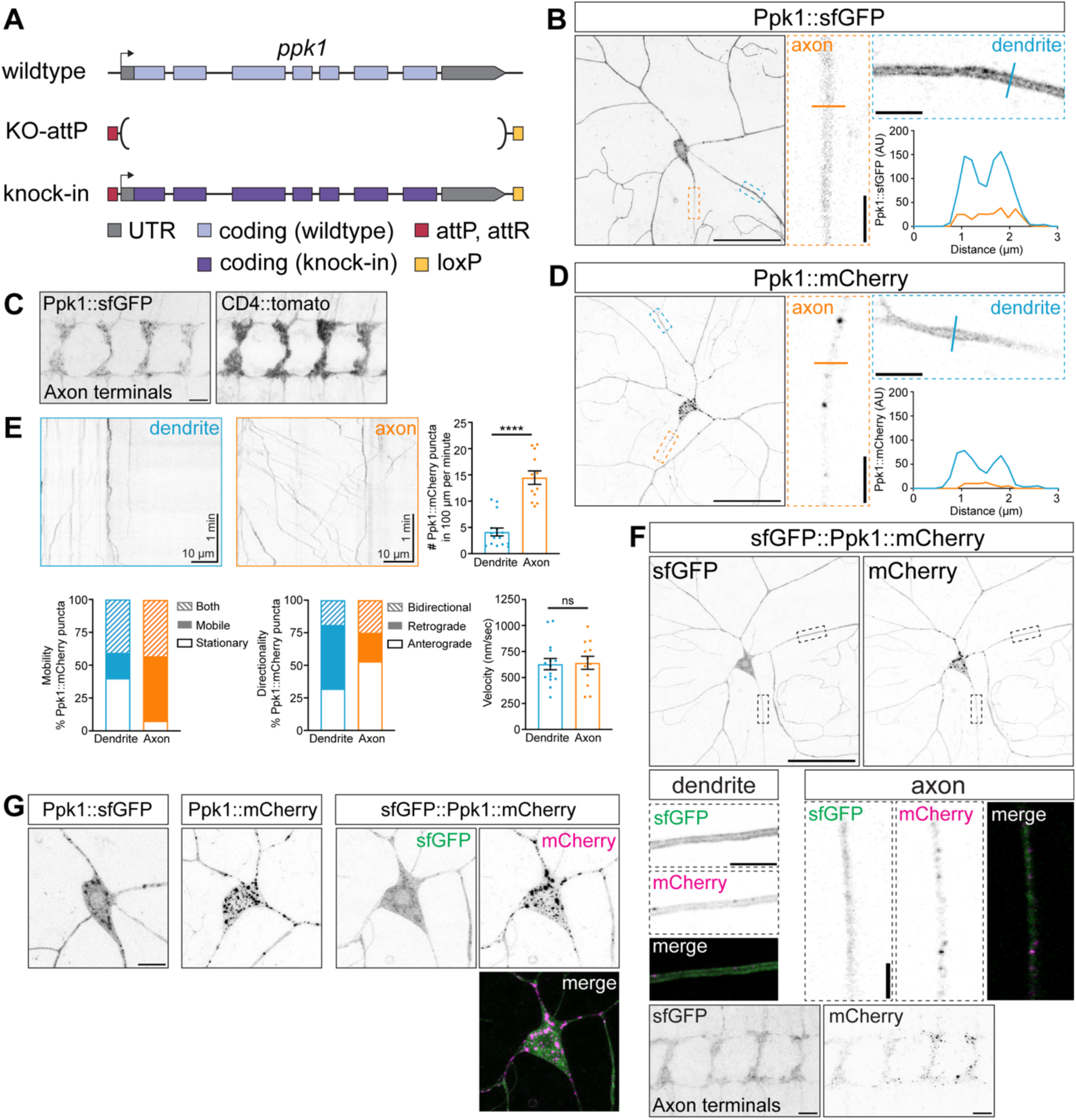
Endogenously tagged Ppk1 is enriched in dendrites but also localizes to axons. Images of ddaC neurons and their axon terminals in live 3^rd^ instar larvae (B, D-F) and fixed ventral nerve cords (VNCs) (C, F). Dashed-outline boxes: Individual 1-µm thick z-plane zoomed-in views of dendrites and axons; a line indicates the position at which an intensity profile plot was generated (B, D). (A) Cartoon illustrating the CRISPR-Cas9-engineered *ppk1* locus in which endogenous *ppk1* was replaced with an attP “docking” site for reliable, rapid knock-in of new *ppk1* alleles. (B, C) Representative images of Ppk1::sfGFP in the cell body, dendrites, and axon (B) and axon terminals in the VNC (C). CD4::tdTomato marks the axon terminals of the class IV neurons in the VNC (C). Scale bars, 50 µm (B), 10 µm (C), and 5 µm (dashed-outline boxes, B). (D) Representative image of Ppk1::mCherry. Scale bars, 50 µm and 5 µm (dashed-outline boxes). (E) Representative kymographs and quantification of Ppk1::mCherry puncta in dendrites and axons. Quantification, Ppk1::mCherry puncta density: Mann-Whitney test (p<0.0001); dendrites (16 dendrite segments, 12 larvae) v. axons (12 axon segments, 9 larvae). In the graph, each data point represents a dendrite or axon segment. Quantification, Ppk1::mCherry puncta motility and directionality: dendrites (16 dendrite segments, 12 larvae) and axons (12 axon segments, 9 larvae). Quantification, Ppk1::mCherry puncta velocity: Student’s unpaired t-test (p=0.8786); dendrites (15 dendrite segments, 11 larvae) v. axons (12 axon segments, 9 larvae). In the graph, each data point represents the average velocity of Ppk1::mCherry puncta in a dendrite or axon segment. Data are plotted as mean ± SEM. n.s.=not significant (p>0.05) and ****p<0.0001. (F) Representative images of dual-tagged sfGFP::Ppk1::mCherry in dendrites, cell body and axon (top and middle), and axon terminal in the VNC (bottom). Scale bars, 50 µm (top), 5 µm (middle), 10 µm (bottom). (G) Representative images of the cell body and proximal dendrites of Ppk1::sfGFP, Ppk1::mCherry, and sfGFP::Ppk1::mCherry. Scale bar, 10 µm.

In addition to tagging Ppk1 with sfGFP, we also tagged Ppk1 with mCherry at the C-terminus. The distribution of Ppk1::mCherry was similar to Ppk1::sfGFP in dendrites and axons (Fig 1D). There was, however, a striking difference: unlike Ppk1::sfGFP, Ppk1::mCherry clustered in bright puncta that concentrated in the cell body, proximal dendrites, and axon (Fig 1D and 1G). These Ppk1::mCherry puncta were motile and moved at speeds consistent with microtubule-based transport (Fig 1E). This is in contrast with Ppk1::sfGFP, which was rarely visible in puncta (Fig 1B). Interestingly, the Ppk1 partner subunit Ppk26 tagged with the fluorescent protein Dendra2 also displays a punctate distribution pattern similar to mCherry-tagged Ppk1, whereas Ppk26 tagged with GFP resembles the more uniform distribution of GFP-tagged Ppk1 [23,28]. Unsure of whether the mCherry tag might direct Ppk1 to a different compartment than sfGFP, we next generated Ppk1 tagged with both sfGFP and mCherry (sfGFP::Ppk1::mCherry). The sfGFP and mCherry fluorescent signals of the dual-tagged sfGFP::Ppk1::mCherry resembled those of the singly tagged Ppk1 proteins (Fig 1F and 1G). This suggests that the punctate mCherry signal does not reflect a difference in the localization of mCherry-tagged Ppk1. Rather, the mCherry tag may reveal the localization of Ppk1 to a compartment(s) where sfGFP does not fluoresce. For example, among other differences, sfGFP and mCherry have different maturation kinetics and are differentially sensitive to pH (e.g., GFP fluorescence is quenched by low pH whereas mCherry is not) [29]. Indeed, these differences in fluorescent protein properties have been leveraged to monitor the subcellular distribution of the transmembrane protein Notch [30]. Our comparison of the fluorescent sfGFP and mCherry signals of the dual-tagged Ppk1 indicates that the identity of the fluorescent protein tag can influence protein visualization and that using different fluorescent tags may be necessary to visualize the full potential range of a protein’s localization in cells.

The localization of fluorescently tagged Ppk1 to both axons and dendrites raises the question of where Ppk1 is inserted into the membrane. To monitor the membrane localization of Ppk1, we initially tagged Ppk1 with superecliptic pHluorin. Superecliptic pHluorin (referred to simply as pHluorin hereafter) is a pH-sensitive GFP variant that is often used to monitor the insertion of transmembrane proteins because it fluoresces in neutral pH environments, such as extracellular space, but has minimal fluorescence in low pH environments, such as the lumen of a transport vesicle [31]. To determine an optimal extracellular position at which to add a fluorescent tag such as pHluorin, we first tagged an extracellular loop of endogenous Ppk1 with sfGFP and compared its fluorescence to Ppk1 tagged with sfGFP at the N- or C-terminus. We tested two positions in an extracellular (EC) loop: Site 1 was selected based on the structure of chicken ASIC1, and Site 2 is a position that was previously used to monitor rat ASIC1a membrane insertion via a haemagglutinin epitope tag [32] (S2 Fig). We found that insertion of sfGFP at Site 1 resulted in fluorescence similar to Ppk1::sfGFP, whereas the insertion of sfGFP at Site 2 led to relatively weak fluorescence (S2 Fig). We then tagged Ppk1 with pHluorin at Site 1 and found that Ppk1::pHluorin^EC^ produced relatively weak fluorescence (S2 Fig). We next tested the effects of eliminating Ppk26 on Ppk1::pHluorin^EC^ fluorescence and distribution, as the loss of Ppk26 should interfere with the membrane localization of Ppk1. In *ppk26* mutant neurons, Ppk1::pHluorin^EC^ fluorescence noticeably increased in the cell body and decreased in dendrites, but fluorescence was still clearly visible in patches in dendrites, particularly at dendrite branch points (S2 Fig). Given that the membrane localization of Ppk1 depends on Ppk26, the fluorescence pattern of Ppk1::pHluorin^EC^ in the *Ppk26* mutant neurons suggested that Ppk1::pHluorin^EC^ might fluoresce in the neutral environment of the endoplasmic reticulum (ER) as well as in the cell membrane. The potential for Ppk1::pHluorin^EC^ to fluoresce in both the ER and cell membrane limits its use as a tool to monitor the insertion of Ppk1 into the neuronal membrane.

The weak fluorescent signal of Ppk1::pHluorin^EC^, and its ability to fluoresce in the ER, led us to develop a new split-GFP approach to monitor the membrane insertion of endogenous Ppk1 (Fig 2A). This approach is analogous to other recently reported split-GFP techniques to monitor the localization of transmembrane proteins [33,34]. We tagged endogenous Ppk1 at Site 1 in the extracellular loop with three copies of the split-GFP peptide GFP(11). In addition to the extracellular GFP(11) tag, we also tagged the Ppk1 C-terminus with mCherry, which enabled us to follow Ppk1 localization throughout the neuron (Ppk1::GFP(11×3)^EC^::mCherry^C-term^). We then expressed a secreted version of GFP(1-10) (secGFP(1-10)) in fat cells, which released secGFP(1-10) into the hemolymph of the larval open circulatory system. Thus, Ppk1 tagged with GFP(11) in an extracellular loop will only fluoresce when Ppk1 is inserted into the cell membrane and encounters secGFP(1-10) (S3 Fig). As a control, we used a construct in which the extracellular N-terminus of the single-pass transmembrane protein CD4 is tagged with GFP(11) and the C-terminus is tagged with tdTomato (GFP(11)^EC^::CD4::tdTomato) (S4 Fig) [35].

**Fig 2.**
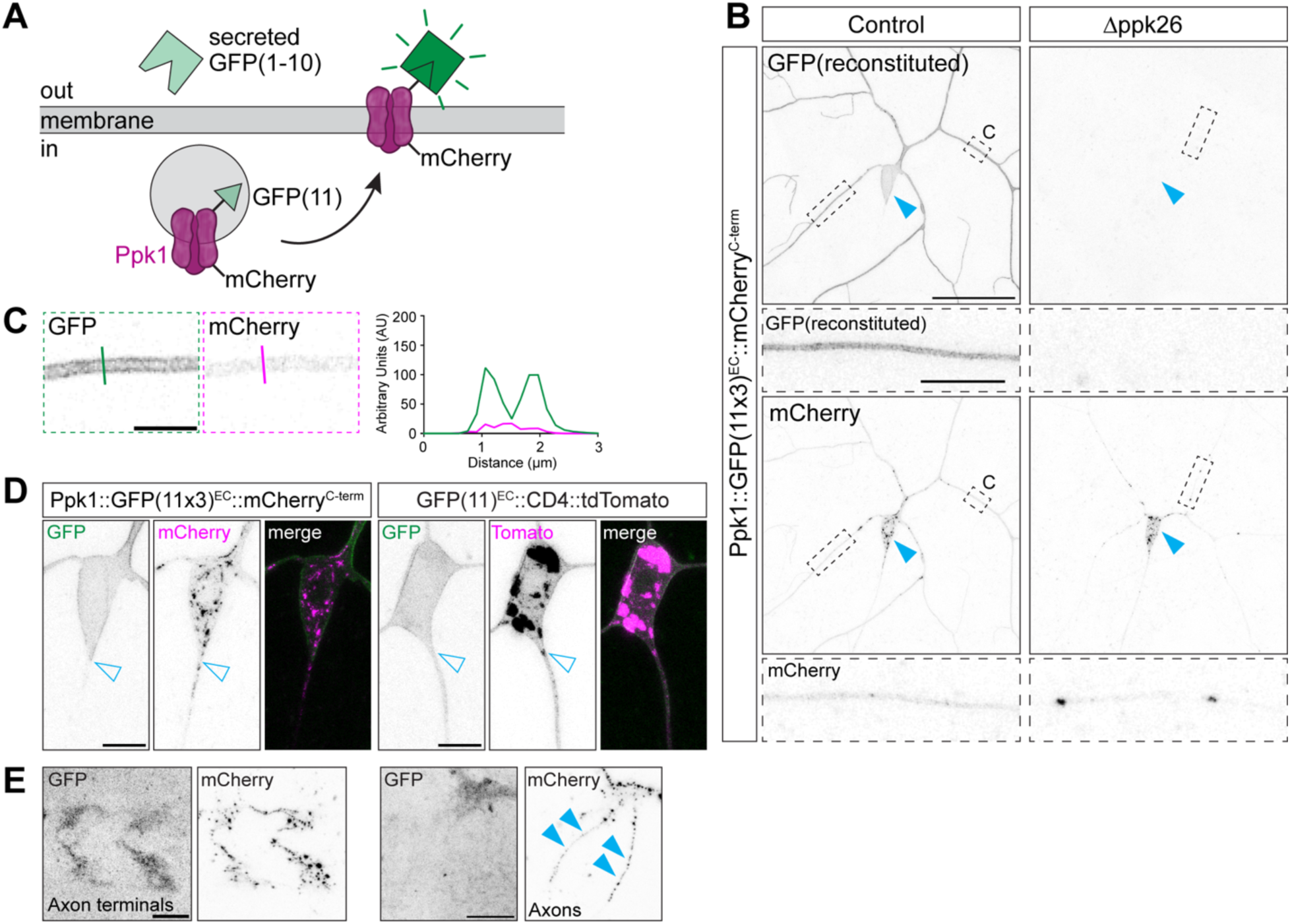
Membrane localization of Ppk1 visualized in live larvae with the secreted split-GFP system. Images of ddaC neurons in live 3^rd^ instar larvae (B-D) and fixed ventral nerve cords (VNCs) (E). Dashed-outline boxes: zoomed-in views of dendrite branches (B, C) and cell bodies (D) from the control neuron shown in panel B. (A) Cartoon of secreted-split-GFP approach to label Ppk1 when it is inserted into the neuronal membrane. Secreted GFP(1-10) (secGFP(1-10)) is expressed by the fat bodies, which secrete secGFP(1-10) into the hemolymph that circulates throughout the larva. When secGFP(1-10) encounters and binds GFP(11) incorporated into an extracellular loop of ppk1, GFP is reconstituted and fluoresces. (B-D) Representative images of GFP and mCherry fluorescence in neurons expressing either Ppk1 dual-tagged with 3 copies of GFP(11) extracellularly (EC) and mCherry at the C-terminus (Ppk1::GFP(11×3)^EC^::mCherry^C-term^) (B-D) or a control construct comprised of the single-pass transmembrane protein CD4 tagged extracellularly with one copy of GFP(11) and intracellularly with tdTomato (GFP(11)^EC^::CD4::tdTomato) (D). UAS-secGFP(1-10) expression was driven in the fat bodies by DcG-Gal4. GFP fluorescence reveals that ppk1 is present in the neuronal membrane of the dendrites (B, C) and cell body (B, D) but not the axon, even though mCherry reveals that Ppk1 is present in axons (B, D). In contrast, GFP(11)^EC^::CD4::tdTomato displays reconstituted GFP signal in both the cell body and the axon (D). Loss of *ppk26* (Δppk26: *ppk26^Δ11/Δ11^*) results in an absence of GFP fluorescence from Ppk1::GFP(11×3)^EC^::mCherry^C-term^ (B). Panel C shows a single 1-µm thick z-plane image of a dendrite segment from the neuron in panel B; the line indicates the position at which the intensity profile plot was generated. AU: arbitrary units. Solid arrowheads: cell body (B); open arrowheads: cell body-axon boundary (D). Scale bars, 50 µm (B, solid-outline boxes), 10 µm (B, dashed-outline boxes; D), and 5 µm (C). (E) Representative images of Ppk1::GFP(11×3)^EC^::mCherry^C-term^ in axon terminals in the VNC. Scale bars, 10 µm. Solid arrowheads: mCherry signal in the axons terminating in the VNC.

Using this split-GFP approach, we monitored GFP fluorescence from Ppk1 and CD4 tagged with GFP(11). Both Ppk1::GFP(11×3)^EC^::mCherry^C-term^ and GFP(11)^EC^::CD4::tdTomato exhibited GFP fluorescence in dendrites and cell bodies (Fig 2B, 2C and 2D, and S4 Fig). As described above, the membrane localization of Ppk1 depends on Ppk26 [21–23]. In neurons lacking Ppk26, there was no GFP fluorescent signal from Ppk1::GFP(11×3)^EC^::mCherry^C-term^, albeit the mCherry signal was still visible, indicating that Ppk1 was still produced in the absence of Ppk26 (Fig 2B). These results support the idea that GFP fluorescence results from Ppk1 insertion into the dendritic membrane.

Next, we examined axons and axon terminals. Strikingly, in neurons expressing Ppk1 tagged with GFP(11), we observed a distinct boundary of GFP fluorescence between the cell body and proximal axon, the latter of which was devoid of fluorescent signal (Fig 2D). While the proximal axon and axon shaft lacked GFP fluorescence, we observed GFP fluorescence in the axon terminals of neurons expressing Ppk1::GFP(11×3)^EC^::mCherry^C-term^ (Fig 2E). The peripheral axons and ventral nerve cord are wrapped by glia, which form a barrier that is unlikely to be penetrated by secGFP(1-10) [36–38]. Thus, the GFP fluorescence from Ppk1 tagged with GFP(11) potentially reflects the trans-endocytosis of Ppk1 from the somatodendritic compartment to the axon terminals.

In contrast to Ppk1 tagged with GFP(11), CD4 tagged with GFP(11) displayed GFP fluorescence throughout the proximal axon and axon shaft (Fig 2D). This indicates a difference in the membrane localization of endogenous Ppk1 and CD4. In summary, we have developed a new assay to monitor the neuronal membrane localization of Ppk1, which we have used to reveal that Ppk1 is inserted into the somatodendritic membrane and the axon terminal membrane but not the membrane in the proximal axon or axon shaft.

### Ppk1 is present in dendritic branches as they form and extend

The localization of Ppk1 and Ppk channels throughout the dendritic arbor raises the question of how Ppk1 becomes so broadly localized. One possibility is that Ppk1 is delivered to dendrites at the same time that the arbor is established, and thus Ppk1 distribution may be coordinated with dendrite growth. To determine the localization of Ppk1 relative to when they develop, we examined the early expression of Ppk1. *ppk1* expression is reported to initiate during embryonic stages, slightly before dendrites emerge (∼embryonic stage 16) [16,21,24–26]. Indeed, we first observed Ppk1::sfGFP in cell bodies and proximal axons just prior to dendrite extension, and we even observe Ppk1::sfGFP in newly emerging dendrites (Fig 3A). These data indicate that Ppk1 is expressed from the very beginning of dendritogenesis.

**Fig 3.**
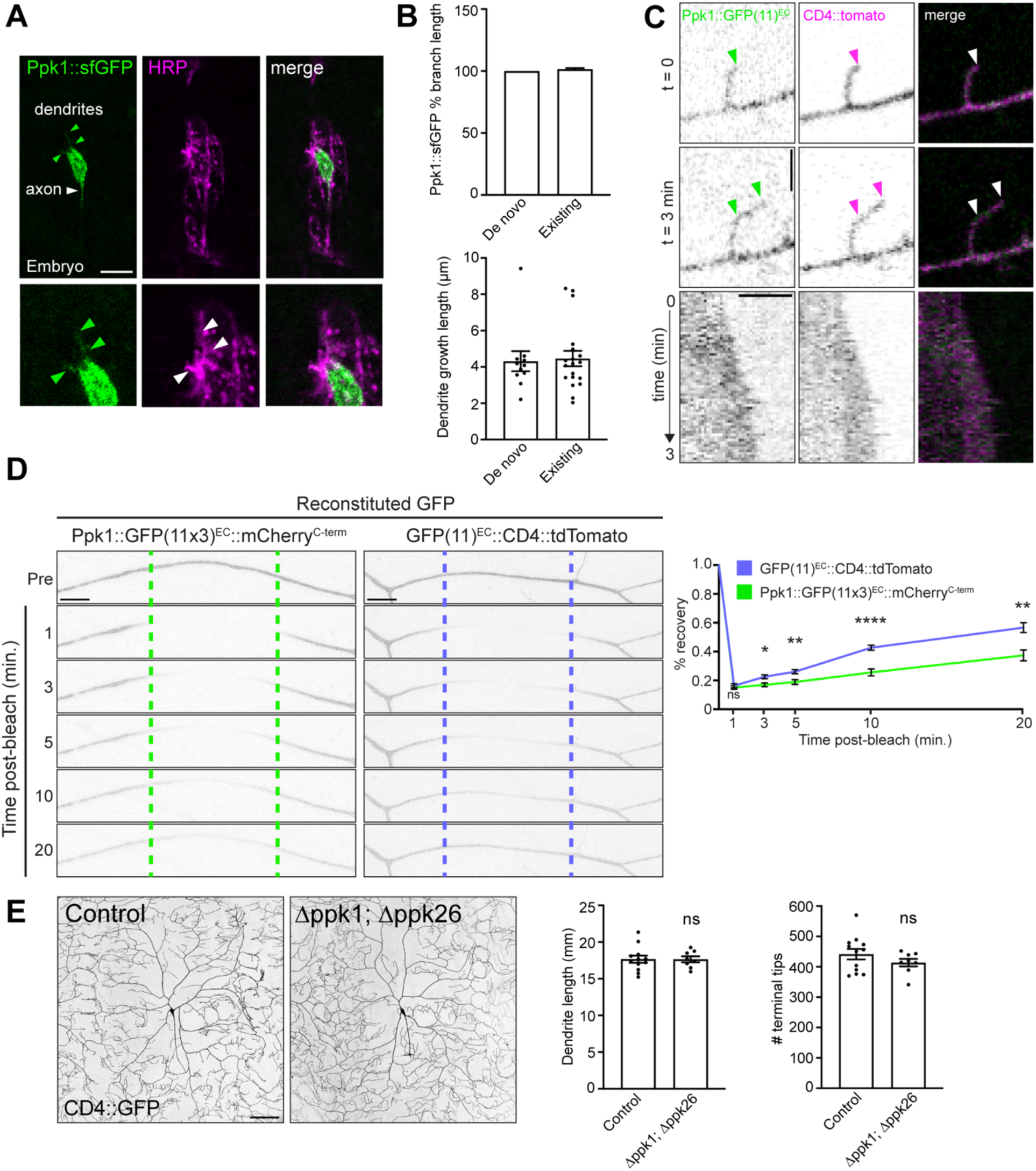
Ppk1 is present in actively growing dendrites but the Pickpocket channel is not required for dendrite growth. Images of ddaC neurons in fixed embryos (A) and live 3^rd^ instar larvae (B-E). (A) Ppk1::sfGFP in the cell body and axon of a ddaC neuron in a late-stage embryo. The ddaC neuron is part of a cluster of sensory neurons, which are marked by anti-HRP. Arrowheads point to Ppk1::sfGFP in nascent dendrites. Scale bar, 10 µm. (B) Quantification of the distribution of Ppk1::sfGFP in growing dendrite branches over 30 minutes. Top graph: In growing branches, Ppk1::sfGFP signal was quantified as a percentage of the branch length marked by CD4::tdTomato (de novo: 11 branches, 4 neurons; existing: 18 branches, 5 neurons). Ppk1::sfGFP was present along the entire length of the branch. Bottom graph: In 30 minutes, both newly formed and existing branches grew equivalent lengths (de novo: 11 branches, 4 neurons; existing: 19 branches, 5 neurons). (C) Still images (top) and kymographs (bottom) from a 3-minute movie of GFP fluorescence from Ppk1::GFP(11×3)^EC^ in a growing dendrite tip marked by CD4::tdTomato. Arrowheads point to initial and final locations of dendrite growth. Genotype: *Ppk1::GFP(11×3)^EC^/DcG-Gal4; UAS-secGFP(1-10)*/*ppk-CD4::tdTomato*. Scale bars, 3 µm. (D) Representative images and quantification of FRAP of GFP in neurons expressing Ppk1::GFP(11×3)^EC^::mCherry^C-term^ and GFP(11)^EC^::CD4::tdTomato (both genotypes: 8 neurons, 8 larvae). Quantification, percent recovery: Student’s unpaired t-test, 1 min (p=0.5417), 3 min (p=0.0108), 5 min (p=0.0064); 10 min (p<0.0001); 20 min (p=0.0021). In the image montage, dashed lines represent the bleached region, which was within 50-150 µm of the cell body. The GFP signal in a 10 µm section centered in the bleached region was quantified at each of the indicated time points. Scale bar, 10 µm. (E) Representative images and quantification of control neurons (*w^1118^*; 12 neurons, 5 larvae) and neurons lacking both Ppk1 and Ppk26 (Δppk1 [*ppk1^attP-KO/attP-KO^*]; Δppk26 [*ppk26^Δ11/Δ11^*]; 8 neurons, 4 larvae). Quantification, dendrite length: Student’s unpaired t-test (p=0.9783). Quantification, dendrite tip number: Student’s unpaired t-test (p=0.2592). The neuronal membrane is marked by CD4::GFP (*ppk-CD4::GFP*). Scale bar, 100 µm. In the graphs, each data point represents either a dendrite branch (B) or a neuron (E), and data are plotted as mean ± SEM. n.s.=not significant (p>0.05), *p<0.05, **p=0.01-0.001, and ****p<0.0001.

Next, we asked whether Ppk1 is present in growing dendrites. Since imaging Ppk1::sfGFP during embryonic stages was technically challenging, we followed the real-time localization of Ppk1::sfGFP in both newly formed and extending dendrite branches in larvae. Strikingly, Ppk1::sfGFP was present throughout growing dendrite branches, in both branches that formed *de novo* and branches that extended (Fig 3B). We imaged Ppk1::sfGFP both continuously and over a period of several minutes, and at both time scales we consistently observed Ppk1::sfGFP throughout dendrite branches as they grew and, in some instances, retracted. We also imaged Ppk1 tagged with GFP(11) and observed GFP fluorescence in growing dendrite branches, which indicates that Ppk1 is integral to the membrane of growing dendrite tips (Fig 3C). Altogether, these data indicate that Ppk1 is present in growing dendrites.

We next asked whether the broad distribution of Ppk1 throughout the dendritic arbor and in growing dendrites might reflect its ability to readily diffuse in the dendritic membrane. To test this we carried out fluorescence recovery after photobleaching (FRAP) with GFP(11)-tagged Ppk1 and CD4 (Ppk1::GFP(11×3)^EC^::mCherry^C-term^ and GFP(11)^EC^::CD4::tdTomato, respectively). After photobleaching, GFP fluorescence from Ppk1::GFP(11×3)^EC^::mCherry^C-term^ recovered gradually over tens of minutes, significantly slower than GFP(11)^EC^::CD4::tdTomato (Fig 3D). These FRAP results indicate that rapid diffusion of Ppk1 is unlikely to explain its broad distribution and presence in growing dendrites.

Our findings that Ppk1 is present in growing dendrites raises the question of whether Ppk1, and its partner subunit Ppk26, are necessary for dendrite growth and whether any change in Ppk channel activity might perturb dendrite morphogenesis. In neurons lacking both Ppk1 and Ppk26, dendrite growth was normal (Fig 3E). Although the Ppk channel is not essential for dendrite growth, we found, consistent with previous reports, that dendrite arborization is reduced when Ppk channel activity is altered by a degenerin mutation in Ppk26, which keeps the channel in an aberrant open state (S5 Fig) [18,21,39,40]. Surprisingly, the loss of Ppk1 enhanced, rather than suppressed, the reduction in dendrite arborization associated with the mutant Ppk26 (S5 Fig). This suggests that the Ppk26 degenerin mutant does not depend on Ppk1 to form a functional channel and that Ppk1 may restrain the activity of the mutant Ppk26.

Our data also suggest that either the mutant Ppk26 may be able to reach the dendritic membrane independently of Ppk1 or that the dendrite growth defect caused by the Ppk26 degenerin mutant is due to the aberrant activity of Ppk26 in an internal organelle. Combined, our results indicate that while Ppk1 and Ppk26 are dispensable for normal dendrite growth, Ppk channel activity must be tightly regulated to achieve proper dendrite arborization.

### Ppk1 levels are not affected by decreasing dendrite arbor size but are reduced when dendrite length significantly increases

Our data reveal that Ppk1 is present in the membrane of extending dendrite branches. One potential model based on these data is that Ppk1 is transported to dendrites via the membrane that grows the dendritic arbor. Since we were not able to visualize the transport of fluorescently tagged Ppk1 in growing dendrites, we instead considered the predictions of this model. One prediction of the model that Ppk1 is transported to dendrites with the membrane that grows the arbor is that the dendritic levels of Ppk1 should be proportional to the amount of dendritic membrane or, in other words, dendrite size. We tested this by assessing Ppk1 levels when dendrite arbor size was altered. We took advantage of different conditions that are known to reduce or increase dendrite growth, acknowledging that these experiments could only serve as a general test of whether Ppk1 levels are proportional to dendrite arbor size. First, we tested two different conditions that reduce dendrite growth: knock-down of the ribosomal protein Rpl22 and overexpression of the ecdysone receptor (EcR) [41,42]. Under both conditions, Ppk1::sfGFP density in dendrites was similar to controls, supporting the idea that Ppk1 levels were proportional to arbor size when dendrite growth was reduced by these two manipulations (Fig 4A and 4B). We then determined whether Ppk1 levels would change when dendrite growth increased using the overexpression of the actin regulator Rac1 [43] and the overexpression of phosphoinositide 3-kinase (PI3K), which regulates cell growth through the mTOR (mechanistic target of rapamycin) pathway [6]. As previously reported, Rac1 overexpression increased dendrite branch number without affecting dendrite length, and we found that Ppk1::sfGFP levels were not affected (Fig 4C). Overexpression of PI3K increased both dendrite branch number and dendrite length by approximately a third, and, in contrast to the other growth-perturbing conditions, Ppk1::sfGFP levels were reduced by nearly a third (Fig 4D). In this overgrowth condition it is possible that the production of Ppk1::sfGFP is outpaced by dendrite growth. Although this is only a small sampling of the many conditions that affect dendrite growth, our results suggest that the dendritic levels of Ppk1 are not generally altered by perturbing dendrite growth.

**Fig 4.**
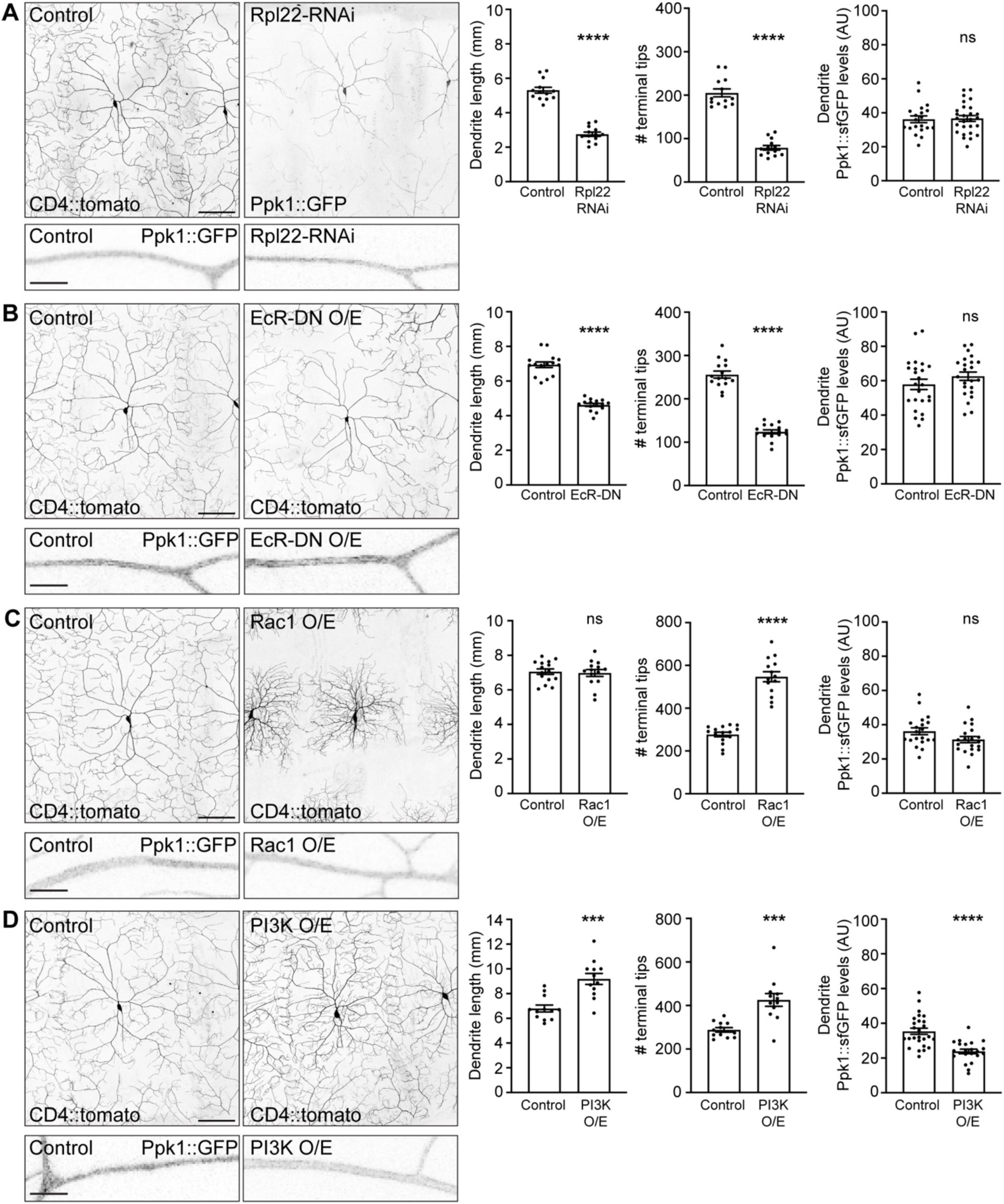
Ppk1 levels are not affected by decreasing dendrite arbor size but are reduced when dendrite length increases. Images of ddaC neuron morphology in live 2^nd^ instar larvae (72 h AEL). The dendritic membrane is marked by CD4::tdTomato, unless otherwise noted. Zoomed inset images show Ppk1::sfGFP in segments of dendrite in live 3^rd^ instar larvae (120 h AEL). (A) Representative images and quantification of control and *Rpl22-RNAi*-expressing neurons. Quantification, dendrite length: Student’s unpaired t-test (p<0.0001); control (13 neurons, 6 larvae) v. *Rpl22-RNAi* (13 neurons, 5 larvae). Dendrite length of *Rpl22-RNAi*-expressing neurons was quantified using Ppk1::sfGFP because the CD4::tdTomato signal was too dim to analyze in these mutants. Quantification, dendrite tip number: Mann-Whitney test (p<0.0001); control (13 neurons, 6 larvae) v. *Rpl22-RNAi* (13 neurons, 5 larvae). Quantification, Ppk1::sfGFP, dendrites: Student’s unpaired t-test (p=0.8602); control (20 neurons, 9 larvae) v. *Rpl22-RNAi* (27 neurons, 14 larvae). (B) Representative images and quantification of control and *EcR-DN*-expressing neurons. Quantification, dendrite length: Student’s unpaired t-test (p<0.0001); control (15 neurons, 9 larvae) v. *EcR-DN* (15 neurons, 7 larvae). Quantification, dendrite tip number: Student’s unpaired t-test (p<0.0001); control (15 neurons, 9 larvae) v. *EcR-DN* (15 neurons, 7 larvae). Quantification, Ppk1::sfGFP, dendrites: Student’s unpaired t-test (p=0.2318); control (24 neurons, 10 larvae) v. *EcR-DN* (23 neurons, 10 larvae). (C) Representative images and quantification of control neurons and neurons over-expressing *Rac1* (*Rac1* O/E). Quantification, dendrite length: Student’s unpaired t-test (p=0.7618); control (15 neurons, 10 larvae) v. *Rac1* O/E (14 neurons, 5 larvae). Quantification, dendrite tip number: Student’s unpaired t-test (p<0.0001); control (15 neurons, 10 larvae) v. *Rac1* O/E (14 neurons, 5 larvae). Quantification, Ppk1::sfGFP, dendrites: Student’s unpaired t-test (p=0.0791); control (20 neurons, 9 larvae) v. *Rac1* O/E (20 neurons, 9 larvae). (D) Representative images and quantification of control neurons and neurons over-expressing *PI3K* (*PI3K* O/E). Quantification, dendrite length: Student’s unpaired t-test (p=0.0001); control (12 neurons, 5 larvae) v. *PI3K* O/E (12 neurons, 6 larvae). Quantification, dendrite tip number: Student’s unpaired t-test (p=0.0002); control (12 neurons, 5 larvae) v. *PI3K* O/E (12 neurons, 6 larvae). Quantification, Ppk1::sfGFP, dendrites: Student’s unpaired t-test (p<0.0001); control (26 neurons, 14 larvae) v. *PI3K* O/E (22 neurons, 13 larvae). Control genotype: *w^1118^*; *ppk-Gal4*. Experimental genotypes: *ppk-Gal4* was used to express the indicated construct. *CD4::tdTomato* and *Ppk1::sfGFP* included as indicated. Experiments to analyze the effects of *Rpl22-RNAi* and *Rac1 over-expression* were run together; the controls for these experiments are the same. In the graphs, each data point represents a neuron, and data are plotted as mean ± SEM. n.s.=not significant (p>0.05), ***p=0.001-0.0001, and ****p<0.0001. AU: arbitrary units. Scale bars, 50 µm.

### Ppk1 persists in dendrites when dynein-mediated transport is perturbed

We further tested the model that Ppk1 is transported to dendrites via the membrane that grows the dendritic arbor by disrupting transport to dendrites. We reasoned that if Ppk1 is trafficked via the membrane that fuels dendrite growth, then perturbing transport to dendrites should decrease growth and have a proportional effect on the dendritic levels of Ppk1 (e.g., the density of Ppk1 should not be affected).

We took several complementary approaches to determine the effects of disrupting dendritic transport on Ppk1. Most transport to and within dendrites occurs along microtubules. In da neuron dendrites, nearly all microtubules are oriented with their minus-ends positioned away from the cell body [44]. Thus, the microtubule minus-end-directed motor dynein is thought to mediate the majority of transport to dendrites. We perturbed dynein function in several ways. First, we reduced the levels of the essential dynein subunit dynein light intermediate chain (Dlic) via RNAi. In *Dlic-RNAi*-expressing neurons, we observed a reduction in dendrite growth as previously reported [45,46], but dendritic Ppk1::sfGFP levels were similar to controls (Fig 5A). Consistent with these results, we found that loss of the dynein co-factor nudE, which also disrupts dendrite morphogenesis [47], did not affect the dendritic levels of Ppk1::sfGFP (control: 112.77 ± 21.05 arbitrary units; *nudE^39A^*/*Df(3L)BSC673*: 110.52 ± 42.55 AU, mean ± standard deviation; Student’s unpaired t-test: p=0.8382; not significant). Thus, decreasing dynein activity reduced dendrite growth, likely reflecting its role in dendritic transport, but Ppk1 levels in dendrites were unchanged. These results indicate that Ppk1 levels remain proportional to dendrite arbor size in dynein loss-of-function neurons.

**Fig 5.**
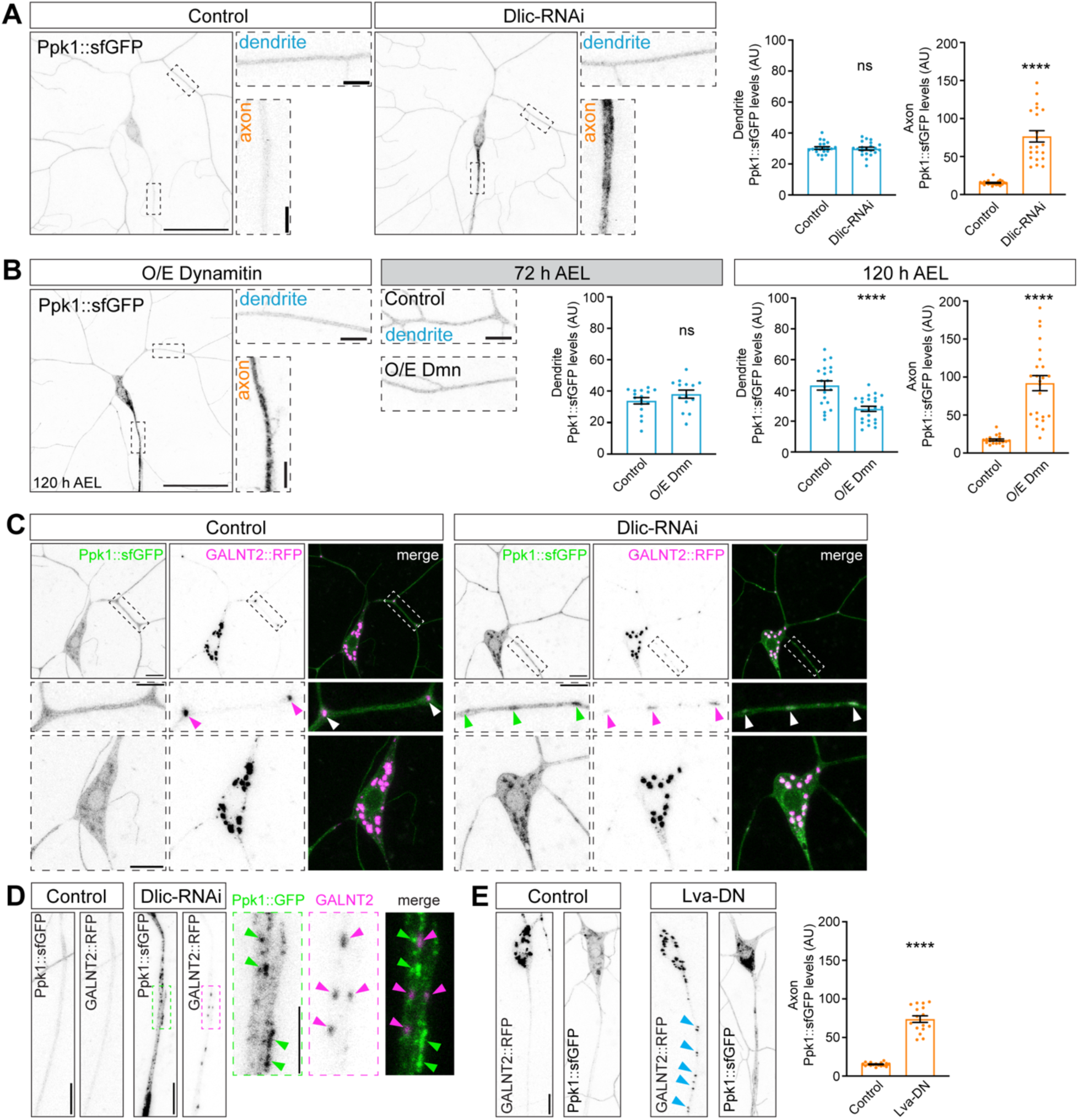
Ppk1 persists in dendrites when dynein-mediated transport is disrupted, and dynein plays a role transporting Ppk1 away from Golgi. Images of ddaC neurons in live 3^rd^ instar larvae. Dashed-outline boxes: zoomed-in views of dendrite branches, cell bodies, and axons. (A) Representative images and quantification of Ppk1::sfGFP in control (21 neurons, 11 larvae) and *Dlic-RNAi*-expressing neurons (21 neurons, 11 larvae). Quantification, dendrites: Student’s unpaired t-test (p=0.7756). Quantification, axons: Mann-Whitney test (p<0.0001). Scale bars, 50 µm and 5 µm (dashed-outline boxes). (B) Representative images and quantification of Ppk1::sfGFP in control neurons and neurons over-expressing *dmn* (O/E *dmn*). Quantification, dendrites, 72 h AEL: Mann Whitney (p=0.1417); control (16 neurons, 6 larvae) v. O/E *dmn* (14 neurons, 6 larvae). Quantification, dendrites, 120 h AEL: Student’s unpaired t-test (p<0.0001); control (20 neurons, 12 larvae) v. O/E *dmn* (24 neurons, 14 larvae). Quantification, axons, 120 h AEL: Mann-Whitney test (p<0.0001); control (20 neurons, 12 larvae) v. O/E *dmn* (23 neurons, 14 larvae). Scale bars, 50 µm and 5 µm (dashed-outline boxes). (C) Representative images of Ppk1::sfGFP and GALNT2::TagRFP in control and *Dlic-RNAi*-expressing neurons. In *Dlic-RNAi*-expressing neurons, Ppk1::sfGFP (green arrowheads) accumulates in Golgi in the cell body and Golgi outposts, marked by GALNT2::TagRFP (magenta arrowheads). Scale bars, 10 µm (solid-outline boxes; dashed-outline boxes, cell body) and 5 µm (dashed-outline boxes, dendrites). (D) Representative images of Ppk1::sfGFP and GALNT2::TagRFP in the axons of control and *Dlic-RNAi*-expressing neurons. Zoomed-in images of an axon of a *Dlic-RNAi*-expressing neuron shows that the Ppk1::sfGFP puncta (green arrowheads) do not colocalize with ectopic Golgi marked by GALNT2::TagRFP (magenta arrowheads). Scale bars, 10 µm and 5 µm (dashed-outline boxes). (E) Representative images of GALNT2::TagRFP and Ppk1::sfGFP in the axons of control and Lva-DN-expressing neurons. In Lva-DN, GALNT2::TagRFP mislocalizes to axons (blue arrowheads) and axonal Ppk1::sfGFP levels increase. Quantification, Ppk1::sfGFP, axons: Student’s unpaired t-test (p<0.0001); control (17 neurons, 6 larvae) v. Lva-DN (16 neurons, 7 larvae). Scale bar, 10 µm. Control genotypes: *w^1118^*; *ppk-Gal4*. Experimental genotypes: *w^1118^*; *ppk-Gal4 UAS-Dlic-RNAi UAS-Dicer* (A, C, D), *w^1118^*; *ppk-Gal4 UAS-dmn* (B), *w^1118^*; *ppk-Gal4 UAS-Lva-DN* (E). *UAS-GALNT2::TagRFP* and *Ppk1::sfGFP* included as indicated (A-E). In the graphs, each data point represents a neuron and data are plotted as mean ± SEM. n.s.=not significant (p>0.05) and ****p<0.0001. AU: arbitrary units.

We also disrupted dynein activity by overexpressing dynamitin (dmn), a dynein co-factor and dynactin complex member. Elevating dmn levels disrupts dynein activity, likely by perturbing dynein-dynactin interactions [48,49]. Unlike *Dlic-RNAi* and *nudE* loss-of-function, the overexpression of *dmn* reduced dendritic levels of Ppk1::sfGFP by approximately a quarter (Fig 5B). Since overexpressed dmn acts as a dominant-negative, it may exert a stronger, cumulative effect on dynein-mediated transport than *Dlic-RNAi* and the *nudE* loss-of-function mutant. Indeed, when we examined younger larvae, neurons overexpressing *dmn* had normal levels of Ppk1::sfGFP (Fig 5B). Such a dose-dependent effect may also explain why we previously observed reduced dendritic Ppk1 levels in clones of *dlic* mutant neurons, which completely lack *dlic* [45]. Our results indicate that the overexpression of *dmn* initially had no effect on Ppk1::sfGFP levels in dendrites but that the persistent overexpression of *dmn* resulted in the decrease of Ppk1::sfGFP dendritic levels over time.

Our findings that the dendritic levels of Ppk1 are not affected by either *Dlic-RNAi* or *nudE* loss-of-function could be interpreted to suggest that Ppk1 is not transported by dynein. To determine whether dynein plays a role in transporting Ppk1 to dendrites, we took a closer look at neurons expressing *Dlic-RNAi*. Most ion channels traffic through the Golgi apparatus, which, in the da neurons, includes both the somatic Golgi and dendritic Golgi “outposts,” which are Golgi mini-stacks found predominantly in the proximal dendritic arbor [50–54]. In control neurons, we found that Ppk1::sfGFP co-localized with somatic Golgi, although we did not detect any Ppk1::sfGFP at dendritic Golgi outposts (Fig 5C). In neurons expressing *Dlic-RNAi*, Ppk1::sfGFP accumulated at somatic Golgi and could also be observed at dendritic Golgi outposts. These results suggest that Ppk1 traffics through both somatic Golgi and Golgi outposts and that Ppk1 is transported away from Golgi by dynein.

In addition to dendritic defects, neurons with altered dynein activity had increased levels of Ppk1::sfGFP in axons (Fig 5A and 5B). Previous work by our group and others has shown that Golgi and Golgi outposts mis-localize to axons when dynein activity is altered [45,47]. Given our findings that Ppk1::sfGFP accumulates at Golgi in dynein loss-of-function neurons, one possibility is that Ppk1::sfGFP “hitchhikes” on Golgi that are ectopically localized to axons. However, in the axons of *Dlic*-*RNAi*-expressing neurons, Ppk1::sfGFP was often adjacent to, but did not overlap with, the trans-Golgi marker GALNT2::TagRFP (Fig 5D). Although Ppk1 was not hitchhiking on Golgi, it was still possible that the mis-localization of Golgi was responsible for the increase in axonal Ppk1 levels. To test this, we perturbed the localization of Golgi by disrupting the golgin lava lamp (lva), which links Golgi to dynein but otherwise does not contribute to dynein activity [55,56]. Consistent with previous reports, we found that the overexpression of dominant-negative lva resulted in the ectopic axonal localization of Golgi [52]. Lva dominant-negative also resulted in an increase in Ppk1::sfGFP levels in axons and the cell body (Fig 5E). Combined, our results suggest that the mis-localization of Golgi in dynein loss-of-function neurons is sufficient to result in the accumulation of Ppk1 in axons.

### Disrupting secretory pathway function impedes the delivery of Ppk1 to the dendritic membrane

Our finding that Ppk1 accumulates at Golgi when dynein function is perturbed indicates that Ppk1 traffics through the Golgi, likely through both the somatic Golgi and Golgi outposts, before being added to the growing dendritic membrane. Based on these data, we hypothesized that interfering with the function of the secretory pathway, which is the predominant source of membrane for dendrite arbor growth and through which membrane proteins traffic, might reduce the dendritic levels of Ppk1. We disrupted secretory pathway function by impeding the budding of cargo-containing vesicles from the ER. We used a transgenic guide RNA (gRNA) that targets *Sec23*, which encodes an essential component of the COPII vesicular coat, in combination with a transgene that expresses Cas9 specifically in the class IV da neurons [57]. The *Sec23* gRNA targets the first coding exon of *Sec23* and, in combination with Cas9, is predicted to generate a loss-of-function allele by introducing a premature stop in Sec23. Indeed, neurons expressing *Sec23* gRNA and Cas9 had shorter dendrites and fewer branches, consistent with previously published work that *Sec23* loss-of-function reduces dendrite morphogenesis (Fig 6A) [52].

**Fig 6.**
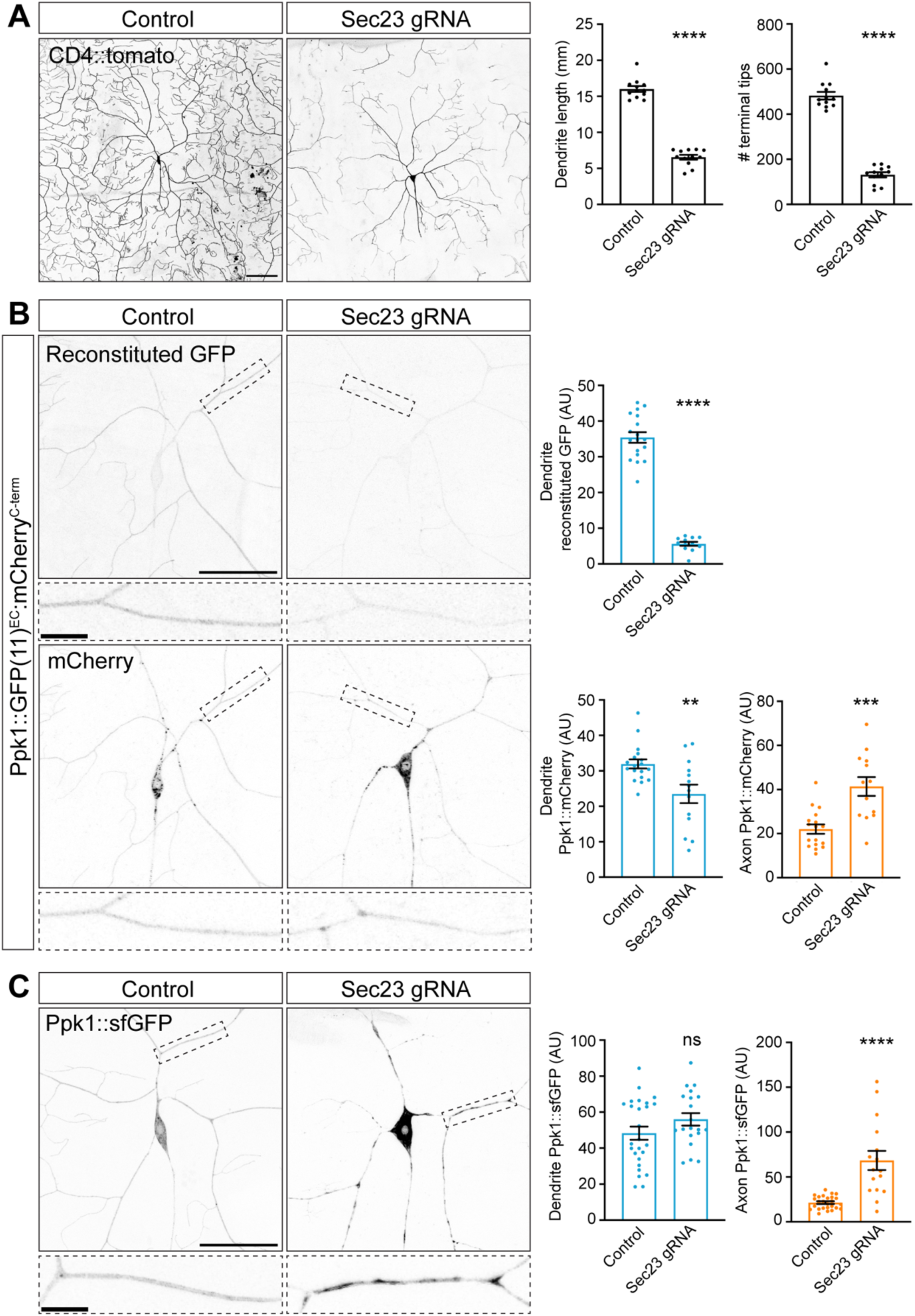
Disrupting secretory pathway function reduces both dendrite growth and membrane-expressed Ppk1. Images of ddaC neurons in live 3^rd^ instar larvae. Dashed-outline boxes: zoomed-in views of dendrite branches. (A) Representative images and quantification of neuron morphology in control (12 neurons, 6 larvae) and *Sec23-gRNA*-expressing (12 neurons, 5 larvae) neurons. The neuronal membrane is marked by CD4::tdTomato. Quantification, dendrite length: Mann-Whitney test (p<0.0001). Quantification, dendrite tip number: Student’s unpaired t-test (p<0.0001). Scale bar, 100 µm. (B) Representative images and quantification of Ppk1::GFP(11×3)^EC^::mCherry^C-term^ in control and *Sec23-gRNA*-expressing neurons. Quantification, reconstituted GFP, dendrites: Student’s unpaired t-test (p<0.0001); control (18 neurons, 8 larvae) v. *Sec23-gRNA* (12 neurons, 8 larvae). Quantification, mCherry, dendrites: Student’s unpaired t-test (p=0.0039); control (18 neurons, 8 larvae) v. *Sec23-gRNA* (14 neurons, 8 larvae). Quantification, mCherry, axons: Student’s unpaired t-test (p=0.0002); control (17 neurons, 8 larvae) v. *Sec23-gRNA* (13 neurons, 8 larvae). Scale bars, 50 µm and 10 µm (dashed-outline boxes). (C) Representative images and quantification of Ppk1::sfGFP in control and *Sec23-gRNA*-expressing neurons. Quantification, dendrites: Student’s unpaired t-test (p=0.1414); control (26 neurons, 15 larvae) v. *Sec23-gRNA* (20 neurons, 11 larvae). Quantification, axons: Student’s unpaired t-test (p<0.0001); control (26 neurons, 15 larvae) v. *Sec23-gRNA* (16 neurons, 10 larvae). Scale bars, 50 µm and 10 µm (dashed-outline boxes). Control genotypes: *w^1118^*; *ppk-Cas9*. Experimental genotype: *w^1118^*; *ppk-Cas9 U6-Sec23-gRNA*. *ppk-CD4::tdTomato*, *Ppk1::GFP(11×3)^EC^::mCherry^C-term^*, and *Ppk1::sfGFP* included as indicated (A-C). In the graphs, each data point represents a neuron, and data are plotted as mean ± SEM. n.s.=not significant (p>0.05), **p=0.01-0.001, ***p=0.001-0.0001, and ****p<0.0001. AU: arbitrary units.

Next, we examined the effects of *Sec23* loss-of-function on Ppk1. *Sec23* loss-of-function neurons had significantly less membrane-associated Ppk1 as revealed by a reduction in GFP fluorescence from the dual-tagged Ppk1::GFP(11×3)^EC^::mCherry^C-term^ (Fig 6B). The mCherry fluorescence of Ppk1::GFP(11×3)^EC^::mCherry^C-term^ was also reduced, but to a lesser extent than the GFP fluorescence. This suggests that disrupting *Sec23* has only a modest effect on Ppk1 levels but has a severe effect on the trafficking of Ppk1 to the neuronal membrane. Consistent with this idea, the dendritic levels of Ppk1::sfGFP were not affected in *Sec23* loss-of-function neurons, although the pattern of Ppk1::sfGFP appeared patchy compared to its typical uniform distribution in control neurons (Fig 6C). Notably, both the mCherry fluorescence of Ppk1::GFP(11×3)^EC^::mCherry^C-term^ and the Ppk1::sfGFP fluorescence appeared greatly increased in the cell bodies (Fig 6B and 6C). Combined, these results suggest that perturbing *Sec23* results in dramatically less membrane-associated Ppk1 and likely causes an accumulation of Ppk1 at an early point during the secretory pathway, likely the ER.

### Dendritic levels of Ppk1 are reduced by disrupting Rab11

We next asked what transport carriers might take Ppk1 from Golgi to growing dendrites and play a role in coordinating ion channel delivery with dendrite growth. We considered Rab11-positive endosomes for several reasons. First, Rab11 has been implicated in the anterograde trafficking of ion channels to the dendritic membrane in mammalian neurons [58], and recent work suggests that Rab11 may affect the trafficking of the Ppk1 partner subunit Ppk26 in fly da neurons [28]. Additionally, Rab11 plays a role in the membrane localization of ENaC family members in the epithelial cells of mammalian kidneys [59–61]. Rab11 mutants also reduce dendrite arborization, indicating that it has a role in dendrite growth (S6 Fig) [28,62]. These data suggest that Rab11 may play a role in both the dendritic localization of Ppk1 and dendrite arbor development.

To test whether Rab11 indeed participates in trafficking Ppk1 to dendrites, we perturbed Rab11 function using both *Rab11-RNAi* and a dominant-negative Rab11 construct, Rab11-DN (Rab11-DN carries an S25N mutation that disrupts GTPase activity). Knocking-down Rab11 with RNAi decreased dendritic Ppk1 as determined by measuring Ppk1 membrane levels via immunohistochemistry, Ppk1::sfGFP, and Ppk1::mCherry (Ppk1 membrane expression was assayed with antibodies rather than GFP(11)-tagged Ppk1 for technical reasons; Fig 7A, 7B and 7C). In neurons expressing *Rab11-RNAi*, the decrease in Ppk1 in dendrites was accompanied by an accumulation of Ppk1::sfGFP and Ppk1::mCherry in cell bodies (Fig 7B and 7C). Similar to *Rab11-RNAi*, the overexpression of Rab11-DN also reduced Ppk1 levels in dendrites as revealed by quantifying Ppk1 membrane expression via immunohistochemistry and Ppk1::mCherry (Rab11-DN is tagged with GFP, and thus we were not able to quantify Ppk1::GFP; Fig 7D and 7E). Like in neurons expressing *Rab11-RNAi*, Ppk1::mCherry accumulated in the cell body in *Rab11-DN*-expressing neurons (Fig 7E). Together, these results indicate that the disruption of Rab11 leads to a decrease in Ppk1 levels in dendrites and an accumulation of Ppk1 in the cell body.

**Fig 7.**
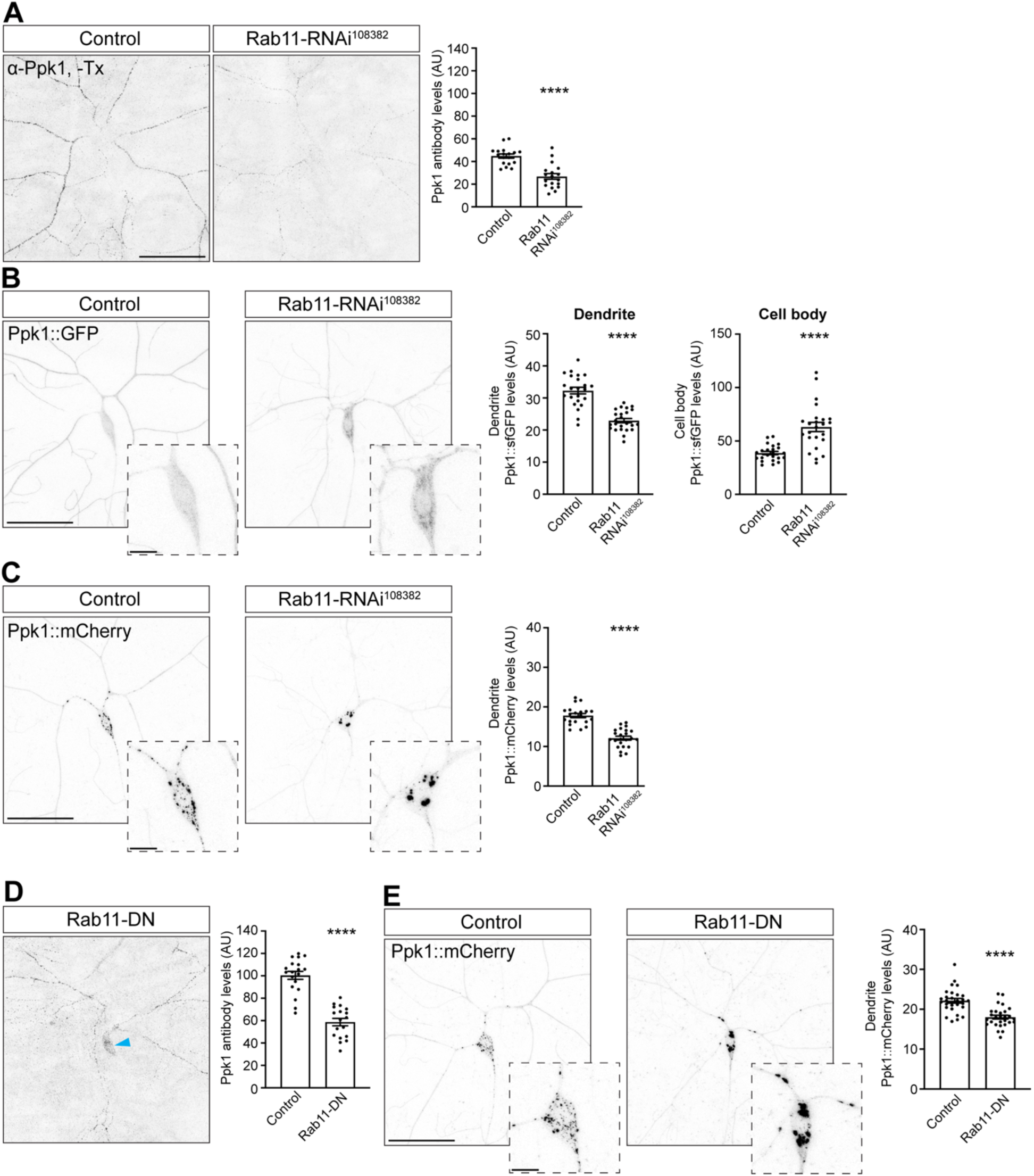
Normal dendritic levels of Ppk1 rely on Rab11. Images of ddaC neurons in 3^rd^ instar larvae, fixed (A, D) or live (B, C, E). Dashed-outline boxes: zoomed-in images of cell bodies. (A) Representative images and quantification of membrane-expressed Ppk1, recognized by anti-Ppk1 antibodies, in control (18 neurons, 6 larvae) and *Rab11-RNAi^108382^*-expressing neurons (18 neurons, 6 larvae). (B) Representative images and quantification of Ppk1::sfGFP in the dendrites and cell bodies of control (23 neurons, 18 larvae) and *Rab11-RNAi^108382^*-expressing neurons (24 neurons, 12 larvae). (C) Representative images and quantification of Ppk1::mCherry in the dendrites of control (20 neurons, 10 larvae) and *Rab11-RNAi^108382^*-expressing neurons (23 neurons, 11 larvae). (D) Representative image and quantification of membrane-expressed Ppk1, recognized by anti-Ppk1 antibodies, in control (19 neurons, 6 larvae) and *Rab11-DN*-expressing neurons (18 neurons, 6 larvae). In ddaC neurons expressing Rab11-DN, Ppk1 immunostaining in the cell body was frequently observed (blue arrowhead). (E) Representative images and quantification of Ppk1::mCherry in control (28 neurons, 12 larvae) and *Rab11-DN*-expressing neurons (28 neurons, 14 larvae). Control genotype: *w^1118^*; *ppk-Gal4*. Experimental genotypes: *w^1118^*; *ppk-Gal4 UAS-Rab11-RNAi^108382^*(A-C) and *w^1118^*; *ppk-Gal4 UAS-Rab11-DN::GFP* (D, E). *Ppk1::sfGFP* or *Ppk1::mCherry* included as indicated (A-E). -Tx: no Triton X-100 (non-permeabilizing conditions). In the graphs, each data point represents a neuron, and data are plotted as mean ± SEM. Quantification: Student’s unpaired t-test. ****p<0.0001. AU: arbitrary units. Scale bars, 50 µm and 10 µm (dashed-outline boxes).

Our findings are consistent with the idea that Rab11 participates in transporting Ppk1 from Golgi to dendrites, but Rab11 also plays a role in the local recycling of ion channels in dendrites [63]. We next asked whether dendritic Ppk1 levels might be regulated via local recycling. In addition to Rab11, which functions in late recycling endosomes, local recycling of ion channels depends on the early endosome GTPase Rab5 [63]. We found that dendritic Ppk1::mCherry puncta colocalized with Rab5-positive early endosomes, as did some Ppk1::mCherry puncta in the cell body, which suggests that Rab5 may play a role in trafficking Ppk1 (Fig 8A).

**Fig 8.**
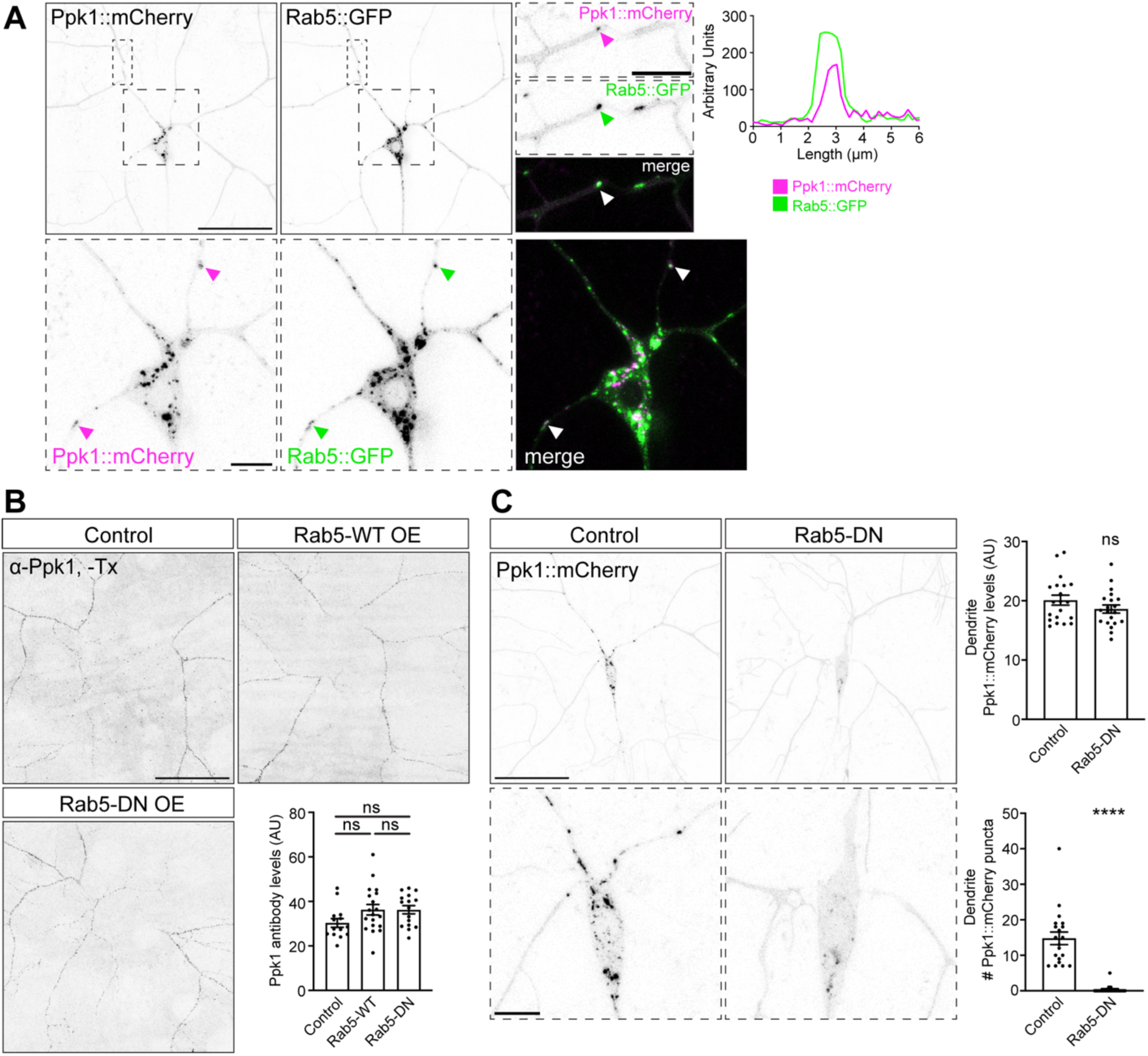
Ppk1::mCherry puncta colocalize with Rab5::GFP, but dendritic levels of Ppk1 do not depend on Rab5 function. Images of ddaC neurons in 3^rd^ instar larvae, live (A, C) or fixed (B). Dashed-outline boxes: zoomed-in views of dendrite branch and cell bodies. (A) Representative images of Ppk1::mCherry with Rab5::GFP. Arrowheads: Colocalized signal. Zoomed-in view of Ppk1::mCherry and Rab5::GFP in a dendrite branch. Line trace is through the punctum highlighted by the arrowhead. Zoomed-in view of Ppk1::mCherry and Rab5::GFP in the cell body and proximal dendrites. (B) Representative images and quantification of membrane-expressed Ppk1, recognized by anti-Ppk1 antibodies, in control neurons (14 neurons, 6 larvae) and neurons over-expressing wild-type *Rab5* (Rab5-WT; 19 neurons, 6 larvae) or *Rab5-DN* (16 neurons, 6 larvae). Quantification: one-way ANOVA with post-hoc Tukey; control v. *Rab5-WT* (p=0.1323), control v. *Rab5-DN* (p=0.1562), and *Rab5-WT* v. *Rab5-DN* (p=0.9999). (C) Representative images and quantification of Ppk1::mCherry in control (20 neurons, 10 larvae) and *Rab5-DN*-expressing neurons (20 neurons, 10 larvae). Quantification, Ppk1::mCherry levels, dendrite: Student’s t-test (p=0.1782). Quantification, Ppk1::mCherry puncta number: Mann-Whitney test (p<0.0001). Ppk1::mCherry puncta were quantified in dendrites within 70 µm of the cell body. Control genotype: *w^1118^*; *ppk-Gal4*. Experimental genotype: *w^1118^*; *ppk-Gal4 UAS-Rab5-DN::YFP* (B, C). *UAS-Rab5::GFP*, *UAS-Rab5::YFP* (Rab5-WT OE), and *Ppk1::mCherry* included as indicated (A-C). -Tx: no Triton X-100 (non-permeabilizing conditions). In the graphs, each data point represents a neuron, and data are plotted as mean ± SEM. n.s.=not significant (p<0.05) and ****p<0.0001. AU: arbitrary units. Scale bars, 50 µm and 10 µm (dashed-outline boxes).

We next analyzed the effects of perturbing Rab5 function on dendritic levels of Ppk1 using a dominant-negative Rab5 construct, Rab5-DN (Rab5-DN carries a S43N mutation that disrupts GTPase activity). The overexpression of Rab5-DN did not disrupt dendritic levels of Ppk1 as measured by quantifying Ppk1 dendritic membrane levels via immunohistochemistry and Ppk1::mCherry (because Rab5-DN is tagged with YFP, we did not quantify GFP-tagged Ppk1) (Fig 8B and 8C). Thus, although Rab5-DN reduces dendrite growth similar to Rab11-DN and *Rab11-RNAi* (S6 Fig and S7 Fig), Rab5-DN has no effect on dendritic Ppk1 levels. Strikingly, although the dendritic levels of Ppk1::mCherry were not affected by Rab5-DN, the number of Ppk1::mCherry puncta in the proximal dendrites and cell body were significantly reduced, and, in many neurons, virtually eliminated (Fig 8C). This suggests that Ppk1::mCherry puncta likely represent Ppk1 in endosomes, specifically early endosomes whose formation depends on Rab5 function. Indeed, consistent with previous reports, we observed that fluorescently tagged Rab5-DN appeared diffuse, which differed from the punctate distribution pattern of fluorescently tagged wild-type Rab5 (S7 Fig) [64]. While this change in Rab5-DN distribution could reflect its dissociation from early endosomes, it might also indicate that early endosome formation is impaired in Rab5-DN-expressing neurons. Overall, our results suggest that Ppk1 traffics through Rab5-positive endosomes but that this trafficking does not have a significant effect on dendritic levels of Ppk1. Combined, our results suggest that Rab11 participates in the forward trafficking of Ppk1 to dendrites at the same time that it promotes dendrite growth.

## Discussion

The perception of stimuli by sensory neurons depends on the morphogenesis of a dendritic arbor equipped with ion channels and receptors that will detect sensory inputs. Central outstanding questions have been: How are ion channels and receptors localized to sensory neuron dendrites during development, and how does a sensory neuron properly match ion channel and receptor levels with dendrite arbor size? Sensory dendrites lack synaptic input, which has made it unclear whether known mechanisms of ion channel and receptor trafficking to dendrites with synapses would also regulate the delivery of ion channels and receptors to the asynaptic dendrites of sensory neurons. Our studies of the localization of Ppk1 suggest a model in which sensory neurons likely package ion channels into the membrane that grows the dendritic arbor, thus coordinating the delivery of ion channels with arbor expansion.

Our data reveal that Ppk1 is present throughout the membrane of dendrites as they grow, indicating that Ppk channels are an integral component of both newly formed and extending dendrites. While the localization and function of ion channels in growing axons is well established [65–71], little is still known about the localization and function of ion channels in developing dendrites. The timing and breadth of Ppk1 distribution indicates that the class IV da sensory dendrites are likely equipped with the capacity to detect stimuli as soon as dendrites emerge. Indeed, recent work has implicated members of the *C. elegans* DEG/ENaC family in sensing mechanical forces to promote terminal branch growth during arbor formation in PVD neurons [72]. Although we found that dendrite development occurred normally without Ppk1 and its partner subunit Ppk26, it is nonetheless possible that Ppk channels participate in dendrite growth, possibly in collaboration with another (mechanosensory) ion channel. Class IV da neurons express at least one additional mechanosensory channel, Piezo, whose worm ortholog is only weakly expressed in PVD neurons. Knockdown of Piezo by RNAi reduces the growth of class IV da neurons [73]. In vertebrate neurons, the distribution of ion channels in young dendrites has been reported but is not well characterized, although a rich body of work supports the role of activity in regulating dendrite growth [74–76]. Our visualization of fluorescently tagged endogenous Ppk1 provides evidence that ion channels are indeed part of the growing dendritic membrane, similar to the localization of ion channels in axons, and that sensory dendrites thus have a “built in” capacity to detect stimuli.

The presence of Ppk1 in growing dendrites raises the possibility that Ppk channels are transported to dendrites as part of the membrane that fuels arbor growth. In one test of this model, we disrupted transport to dendrites by interfering with dynein. Dynein is the predominant motor for dendritic transport in da neurons, and perturbing dynein function significantly reduces arbor size [45,46]. We found that the dendritic levels of Ppk1::sfGFP remained normally proportional to arbor size in neurons with reduced dynein activity. This suggests that in dynein loss-of-function neurons, the membrane that does make it to dendrites has normal levels of Ppk1. The finding that Ppk1::sfGFP accumulates at Golgi in dynein loss-of-function neurons indicates that dynein normally transports Ppk1 (and Ppk channels) from Golgi to dendrites. Our results, combined with previous studies [46], suggest that interfering with dynein function reduces the amount of membrane that is added to the arbor over time but does not affect the amount of Ppk1 that is packaged into the membrane destined for dendrites. Thus, interfering with dynein activity perturbs dendrite arborization but does not affect Ppk1 density. Strikingly, even interfering with translation via the knockdown of the ribosomal protein RpL22, which decreases dendrite growth, does not affect dendritic levels of Ppk1. This result is also consistent with the idea that Ppk1 and Ppk channels are packaged into transport packets at a consistent density even if their subsequent transport out of the Golgi is disrupted. Moreover, our analysis of Ppk1::sfGFP levels in dendrite growth mutants, coupled with published reports of Ppk1 or Ppk26 in mutants that affect arbor size, also indicate that Ppk levels typically remain constant or are only minimally affected despite dramatic changes in dendrite arbor size [77–79]. The idea that Ppk1 and Ppk channels are transported to dendrites via the membrane that grows dendrites suggests a mechanism for the coordination of ion channel levels and dendrite arbor size.

Once Ppk1 reaches the dendrites, our FRAP analysis of membrane-localized Ppk1 (Ppk1::GFP(11)^EC^) suggests that Ppk channels remain stably inserted into the dendritic membrane with little baseline turnover. This idea is also supported by our finding that Ppk1 levels are not significantly affected when local protein recycling is disrupted via the expression of Rab5-DN. Moreover, our FRAP analysis of Ppk1::GFP(11)^EC^ did not uncover any hot spots of fluorescent signal recovery that might indicate areas of ion channel addition, although it is possible that such sites of insertion are below our level of detection. Recent qRT-PCR analysis has revealed that Ppk1 mRNA levels drastically decrease after the 2^nd^ instar developmental stage [26], which corresponds to when dendrite growth slows significantly (Parrish et al. 2009). These data, combined with our FRAP results, suggest that Ppk1 is stably integrated into the dendritic membrane as dendrites grow expansively, and, once dendrite growth plateaus, there is little replenishment with newly expressed Ppk1.

To monitor the membrane localization of Ppk1, we used a novel split-GFP approach. These experiments revealed that Ppk1 is present throughout the somatodendritic membrane but not the membrane of the proximal axon or axon shaft. The clear demarcation of Ppk1::GFP(11)^EC^ fluorescent signal between the somatodendritic and axonal compartments is consistent with the presence of a diffusion barrier between these two compartments in the proximal axon of fly neurons, as was previously supported by studies of the fly ankyrin Ank2 [80–82]. Although there is a barrier to the diffusion of Ppk1 from the somatodendritic membrane to the axonal membrane, Ppk1 is not prevented from being trafficked into the axon to the axon terminal. Given that ensheathing glia likely exclude secGFP(1-10) [36–38], the Ppk1::GFP(11)^EC^ fluorescent signal at axon terminals may reflect the trans-endocytosis of Ppk1 from dendrites to axons. It is not clear what role Ppk1 may play in the axon terminals of class IV da neurons, although other studies, including in flies, support presynaptic roles for Ppk channels and their orthologs [18,19,83]. Additional studies will be needed to determine whether Ppk1 is preferentially trafficked to one compartment or the other and whether Ppk1 may be stably inserted into the dendritic, but not axonal, membrane. Ppk1::GFP(11)^EC^ provides a tool to investigate the spatially restricted membrane expression of Ppk1 and the diffusion barrier between the somatodendritic and axonal membranes.

Our data indicate that Rab11 plays a role in the transport of Ppk1 to dendrites. Rab11 is an integral component of recycling endosomes but has also been implicated in the anterograde trafficking of receptors and ion channels, including ENaCs, in neuronal and non-neuronal cells [58,59,84–86]. Our results implicate Rab11 in the forward transport of Ppk1 in fly da neurons, which points to the potential conservation of DEG/ENaC/ASIC trafficking pathways across organisms and cell types. Our results indicate that disrupting Rab11 reduces Ppk1 in dendrites but that disrupting the function of Rab5, which acts in early endosomes, does not. This suggests that the effects of perturbing Rab11 on Ppk1 is not due to an effect on Rab11-mediated recycling but is likely due to a disruption of Rab11-mediated forward trafficking of Ppk1 to dendrites. Consistent with this model, Ppk1 levels increase in the cell body when Rab11 levels or function are perturbed. Recently, another group has also found that Ppk26 levels also increase in the cell bodies of neurons with altered Rab11 [28]. It is notable that disrupting Rab11 does not lead to a total loss of Ppk1 from dendrites. This may be due to incomplete perturbation of Rab11 or Rab11-positive endosomes, or it could indicate a complementary pathway for the transport of Ppk1 and Ppk channels to dendrites. The pathways that supply membrane and membrane proteins to dendrites, particularly growing dendrites, are still poorly understood. Our data suggest that Rab11 and Rab11-positive endosomes may participate in a pathway that coordinates Ppk ion channel delivery and dendrite arbor expansion.

## Materials and Methods

### Fly husbandry and stocks

Fruit flies were maintained at 25°C on cornmeal-molasses-yeast medium. The generation of new *ppk1* alleles and the *UAS-secGFP(1-10)* flies are described below. *ppk26* strains including *ppk26^Δ11^*, *UAS-ppk26::mCherry*, and *UAS-ppk26-DEG(A547V)::mCherry* were gifts of Dr. Yuh Nung Jan (UCSF) [21]. The following alleles and transgenic fly strains from the Bloomington Drosophila Stock Center (BDSC), Vienna Drosophila Resource Center (VDRC), and individual laboratories were used: *ppk-Cas9* [57], *Df(3L)BSC673* (BDSC 26525), *ppk-CD4::tdTomato* (BDSC 35845), *ppk-GFP(11)^EC^::CD4::tdTomato* [35], *hsp70-Cre* (BDSC 1092), *DcG-Gal4* [87], *UAS-Dcr-2* (BDSC 24650), *UAS-Dlic-RNAi* (VDRC 41686), *UAS-dynamitin* (BDSC 8784), *UAS-EcR-DN* (BDSC 9449), *UAS-GALNT2::TagRFP* (BDSC 65253), *UAS-sfGFP(1-10)* (Bo Huang, UCSF), *UAS-Lva-DN* (BDSC 55055), *nudE^39A^* [88], *Ppk-Gal4* (BDSC 32078, BDSC 32079), *UASp-Rab5-WT::YFP* (BDSC 24616), *UASp-Rab5-DN[S43N]::YFP* (BDSC 9772), *UAS-GFP::Rab5* (BDSC 43336), *UAS-PI3K* (BDSC 8294), *UAS-Rab11-RNAi* (VDRC 108382), *UAS-Rab11-DN(3-4)::GFP* (Hsiu-Hsiang Lee, National Taiwan University College of Medicine), *UAS-Rac1* (BDSC 6293), *UAS-RpL22 RNAi* (BDSC 34828), *U6-Sec23-gRNA* (BDSC 79400), *w^1118^*.

### Generation of *ppk1* alleles

The endogenous *ppk1* gene (∼3.7 kb encompassing the entire *ppk1* transcript) was knocked-out and replaced with an attP site to facilitate the reliable integration of new *ppk1* alleles. The *ppk1^attP-KO^* strain was generated using CRISPR-Cas9 genome engineering and ends-out gene-targeting [89–91]. We used two guide RNAs that flank *ppk1* (upstream gRNA: 5’-GTTCTTATATCTAGAGATGT-3’, and downstream gRNA: 5’-GTCAAGACTTGAAGAATACTG-3’) and a donor template, which contained homology arms surrounding an attP site and 3xP3-DsRed flanked by loxP sites. Candidate strains were identified by DsRed expression in adult eyes, and the *ppk1* locus was analyzed by sequencing genomic DNA from isogenized candidate strains. 3xP3-DsRed was then removed by crossing to flies expressing Cre recombinase. A single strain with the desired replacement of *ppk1* with an attP site was used to generate all knock-in alleles.

Constructs to create knock-in alleles were generated using standard molecular biology techniques and Gibson Assembly to add molecular tags. Two plasmid backbones were used: *pGE-attB-GMR* [90], which includes a *GMR-mini-white^+^* cassette to identify knock-in alleles by red eye color in adults, and *pBSK-attB-3xP3* (this study), which includes a *3xP3-DsRed* cassette to identify knock-in alleles by DsRed expression in adult eyes. *pBSK-attB-3xP3* was generated by adding an *attB* site and *3xP3-DsRed* to *pBSK*. All the exogenous sequences (e.g., the product of attB/attP recombination) knocked-into the endogenous *ppk1* locus were the same regardless of which plasmid backbone was used. New *ppk1* alleles in the *pGE-attB-GMR* vector were first subcloned into *pBSK*, modified, and then inserted into *pGE-attB-GMR* using EcoRI and KpnI. New *ppk1* alleles in the *pBSK-attB-3xP3* plasmid were cloned directly using Gibson Assembly (*pBSK-attB-3xP3-ppk^WT^* was the starting plasmid for many of the knock-in alleles). All constructs were verified by sequencing prior to injection. *attB*-containing plasmids with *ppk1* knock-in alleles were injected into *ppk1^attP-KO^* embryos expressing ΦC31 integrase (BestGene Inc., Chino Hills, CA). The *GMR-mini-w^+^* and *3xP3-DsRed* markers were subsequently removed by crossing to flies expressing Cre recombinase.

The following alleles were generated in this study: First a wild-type knock-in allele (*ppk1^WT-k’in^*) was generated by cloning the part of the *ppk1* locus that was eliminated by the replacement strategy (plasmid *pGE-attB-GMR-ppk1^WT^)*. The resulting *ppk1^WT-K’in^* flies displayed no overt phenotypes and restored the normal pattern of *ppk1* expression. sfGFP::Ppk1 and Ppk1::sfGFP were created by adding one copy of sfGFP and a GGS(x4) linker at the N- or C-terminus, respectively, of Ppk1 (plasmids *pGE-attB-GMR-sfGFP::ppk1* and *pGE-attB-GMR-ppk1::sfGFP*). A similar approach was used to generate Ppk1::mCherry and sfGFP::Ppk1::mCherry (plasmids *pBSK-attB-3xP3-ppk1::mCherry* and *pBSK-attB-3xP3-sfGFP::ppk1::mCherry*). Ppk1::sfGFP^EC^-Site 1 and Ppk1::sfGFP^EC^-Site 2 were created by tagging Ppk1 with one copy of sfGFP at either extracellular Site 1 (between Asn171 and Ile172) or Site 2 (between Gln204 and Leu205). sfGFP was flanked on both sides by a GGS(x4) linker (plasmids *pBSK-attB-3xP3-ppk1::sfGFP^EC^-Site 1* and *pBSK-attB-3xP3-ppk1::sfGFP^EC^-Site 2)*. Ppk1::GFP(11×3)^EC^ was created by tagging Ppk1 with three copies of the split-GFP peptide GFP(11) at extracellular Site 1 (between Asn171 and Ile172). The three copies of GFP(11) were flanked on both sides by a GGS(x4) linker (plasmid *pBSK-attB-3xP3-ppk1::GFP(11×3)^EC^*). Ppk1::GFP(11×3)^EC^::mCherry^C-term^ was created by tagging Ppk1::GFP(11×3)^EC^ at the C-terminus with one copy of mCherry connected by a GGS(x4) linker (plasmid *pBSK-attB-3xP3-ppk1::GFP(11×3)^EC^::mCherry^C-term^*).

Ppk1::GFP(11×7)^C-term^ was created by tagging Ppk1 with seven copies of GFP(11) at the C-terminus via a GGS(x4) linker (plasmid *pGE-attB-GMR-ppk1::GFP(11×7)*). Ppk1::pHluorin^EC^::mScarlet^C-term^ and Ppk1::pHluorin^EC^::mCherry^C-term^ were created by adding one copy of superecliptic pHluorin (synthesized as a gene block by GeneWiz, South Plainfield, NJ) at extracellular Site 1 (between Asn171 and Ile172) and flanked on both sides by a GGS(x4) linker and tagged at the C-terminus with either one copy of mScarlet-I (synthetic gene block from GeneWiz) or one copy of mCherry connected with a GGS(x4) linker (plasmids *pGE-attB-GMR-ppk1::pHluorin^EC^::mScarlet^C-term^*and *pBSK-attB-3xP3-ppk1::pHluorin^EC^::mCherry^C-term^*).

### Generation of the UAS-secGFP(1-10) transgenic fly strain

The GFP(1-10) coding sequence, which was synthesized as a gBlock fragment (Integrated DNA Technologies, Inc.), was PCR-amplified and cloned into the NheI/XbaI sites of *pIHEU-sfGFP-LactC1C2* [92]. The resulting *pIHEU-secGFP(1-10)* construct contains a signal-peptide sequence from Adipokinetic hormone fused in-frame before GFP(1-10). The construct was inserted at the *attP^VK00005^*site (injected by Rainbow Transgenic Flies, Inc).

### Fixation and Immunohistochemistry

To visualize Ppk channel expression in larval fillets, wandering third instar larvae were washed in 1X PBS (phosphate buffered saline, pH 7.4), dissected in PHEM buffer (80 mM PIPES pH 6.9, 25 mM HEPES pH 7.0, 7 mM MgCl_2_, 1 mM EGTA) and fixed in 4% paraformaldehyde in 1X PBS with 3.2% sucrose for 20 minutes. For the dissection, larvae were pinned onto a Sylgard plate with their dorsal trachea facing down and were cut on their ventral side to preserve the ddaC neurons. After fixation, the dissected fillets were washed 3 times with 1X PBS, quenched with 50 mM NH_4_Cl for 10 minutes, and blocked in blocking buffer composed of 2.5% bovine serum albumin (BSA; catalog number A9647, Sigma), 0.25% fish-skin gelatin (FSG; catalog number G7765, Sigma), 10 mM glycine, and 50 mM NH_4_Cl for 3 hours at room temperature. Fillets were incubated in primary antibody diluted in blocking buffer overnight at 4°C. The next day, fillets were washed in 1X PBS at room temperature (3 x 30 minutes) and incubated with secondary antibody diluted in blocking buffer overnight at 4°C. The next day, fillets were washed in 1X PBS at room temperature (3 x 30 minutes) and mounted onto glass microscope slides (Fisher Scientific, Selectfrost, 25×75×1.0 mm) with cover glass (Fisher Scientific 24×50-1.5) using elvanol containing antifade (polyvinyl alcohol, Tris-HCl pH 8.5, glycerol and DABCO, catalog number 11247100, Fisher Scientific, Hampton, NH). All wash and incubation steps were performed on a nutator. To visualize membrane levels of Ppk1, a rabbit anti-Ppk1 antibody (1:3000; gift of Yuh Nung Jan, UCSF) [21] targeting an extracellular epitope of Ppk1 was used without detergent in any wash or incubation steps. A fluorescently conjugated secondary antibody was used: goat anti-rabbit-Dylight 633 (1:500; catalog #35563, Invitrogen).

To visualize Ppk expression in the ventral nerve cord, brains from wandering third instar larvae were isolated from larval carcasses in 1x PBS. Following fixation (4% paraformaldehyde in 1X PBS with 3.2% sucrose for 15 minutes), brains were washed (3 X 5 minutes) in 1x PBS and mounted with the optic lobes facing down. The cover glass was stabilized with four small dots of vacuum grease spacers in four corners of the slide.

To visualize Ppk1::sfGFP in young ddaC neurons, embryos were collected on grape plates for several hours and then devitalized in a solution of 50% bleach and 50% H_2_O for 2-3 minutes. The eggshells were washed away by rinsing with H_2_O and then placed in a tube containing equal quantities of n-heptane and fixative (4% paraformaldehyde in 1X PBS) for 10 minutes. Fixed embryos were washed with 1X PBS with 0.1% Triton X-100 (3 x 10 minutes) and then probed for 2 hours with goat anti-HRP conjugated Alexa Fluor 647 (1:1000, or 0.5 mg/mL, Jackson ImmunoResearch, West Grove, PA). The anti-HRP antibody recognizes a glycoprotein epitope that is present throughout the fruit fly nervous system, enabling visualization of virtually all neuronal membranes. Following incubation with the anti-HRP antibody, the embryos were washed in 1X PBS with 0.1% Triton X-100 (3 x 30 minutes). Embryos were mounted in a solution of 50% glycerol and 50% 1X PBS on glass microscope slides (Fisher Scientific, Selectfrost, 25×75×1.0 mm) with cover glass (Global Scientific, 24×50 mm-1.5). All steps were performed at room temperature.

### Imaging

Imaging was performed on either an SP5 or Stellaris laser-scanning confocal microscope (Leica Microsystems) with sensitive hybrid (HyD) and photomultiplier tube (PMT) detectors using 20×0.7 NA (SP5), 20×0.75 NA (Stellaris), and 40×1.3 NA (SP5 and Stellaris) oil-immersion objectives. The dorsal class IV da neurons (ddaCs) in abdominal segments A2-A5 of control and mutant larvae were imaged. For live imaging, individual larvae were placed into a small drop of 50% glycerol:1X PBS solution that was flanked on both sides by strips of vacuum grease spacers. The larva was then immobilized by pressing a cover glass on top of the spacers. The larva was oriented with its dorsal trachea facing up and rolled gently to one side for optimal positioning of the ddaC neurons. Fixed samples (larval fillets, VNCs) were imaged using a 40×1.3 NA oil-immersion objective. Images were collected via z-stacks (1024×1024-pixel resolution, 1 µm per z-step). Movies of Ppk1::sfGFP and Ppk1::mCherry dynamics were collected in 120 h AEL larvae using a 40×1.3 NA oil-immersion objective at a resolution of 1024 x 256 pixels, zoom 2.5, and a rate of 2-1.35 frames per second, or 0.5-0.74 seconds per frame, respectively, for a duration of 3 minutes. Movies of Ppk1::GFP(11)^EC^ in growing dendrite tips were collected in 72 h AEL larvae using a 40×1.3 NA oil-immersion objective at a resolution of 1024 x 256 pixels, zoom 6, and a rate of 0.34 frames per second (2.942 seconds per frame) for a duration of 3 minutes. For FRAP: First, a pre-bleach z-stack was obtained of the dendrite region to be bleached, which was a secondary dendrite segment longer than 50 µm without branch points, visible in a single z-plane, and within 150 µm of the cell body. A 50 µm circular region of interest (ROI) was centered on the dendrite segment. Next, the Leica FRAP Wizard was used to bleach the ROI: pre-bleach (10 frames; 0.739 sec/frame), bleach at 100% 488 laser intensity (10 frames; 0.739 sec/frame), post-bleach (10 frames; 0.739 sec/frame). After bleaching, z-stacks (z-step size of 0.5 µm) were captured at 1, 3, 5, 10, and 20 minutes post-bleaching. For all experiments, the same imaging settings were used for control and experimental conditions. Images and movies were subsequently analyzed using FIJI or Metamorph.

### Quantification of Ppk1 signal

Levels of Ppk1 were measured in FIJI using the following reporters: anti-Ppk1 antibody, Ppk1::sfGFP, Ppk1::mCherry, and Ppk1::sfGFP(11×3)^EC^::mCherry^C-term^. First, maximum intensity projections of z-stack images were generated. To quantify levels in dendrites, the fluorescence intensity of three different dendrite branches was quantified by tracing 50-µm lines over segments close to the cell body and averaging the signal intensity along each segment. The average intensity of a 50-µm line traced over the background was subtracted from each dendrite trace. Under non-permeabilizing conditions, anti-Ppk1 signal was weak surrounding the cell body, and therefore dendrite traces initiated ∼10-15 µm away from the cell body. An average intensity for each neuron was quantified by averaging the intensities of the three dendrite segments after subtracting the background signal. A similar protocol was used to determine the fluorescence intensity of Ppk1::sfGFP in axons: a 50-µm line was traced over the axon close to the cell body and the average intensity of a 50-µm line traced over the background was subtracted.

To measure the extent of Ppk1::sfGFP signal in growing dendrites over time, z-stack images of Ppk1::sfGFP and CD4::tdTomato taken at two time points 30 sec apart were aligned using the bUnwarpJ plugin to generate a composite image representing the change in dendrite length and fluorescent signal over time. Dendrite length was quantified based on the CD4::tdTomato signal, and the percentage of dendrite that was Ppk1::sfGFP-positive was calculated.

### Quantification of Ppk1::sfGFP and Ppk1::mCherry dynamics

Movies of Ppk1::sfGFP and Ppk1::mCherry dynamics were first stabilized in FIJI using the Image Stabilizer plugin. Stabilized movies were opened in Metamorph (Molecular Devices, LLC, San Jose, CA), and kymographs were generated by drawing 50-70 µm line segments along dendrites and axons. Frequency was quantified by manually counting the number of motile and stationary puncta (data were normalized to represent the number of puncta in 100 µm and 1 minute). To quantify motility, the tracks on each kymograph were manually divided into three categories. Puncta that were motile for the duration of the movie were scored as mobile. Some puncta were both motile and stationary; these puncta were scored as “both.” Puncta that did not move (defined as less than 1 µm) for the duration of the 3-minute movie were scored as stationary. To quantify directionality, the tracks on the kymograph were manually scored as anterograde (away from the cell body), retrograde (towards the cell body), or bidirectional (anterograde and retrograde movement). To quantify velocity, tracks on the kymograph were traced, and the corresponding data on time and distance were exported to Excel to calculate velocity.

### Quantification of dendrite morphology

Imaris software with Filament Tracer (version 9.7-9.8, Oxford Instruments) was used to quantify dendrite length and the number of terminal tips. Neurons were analyzed in larvae that were aged to 72 h AEL unless otherwise mentioned. To capture the entire ddaC dendritic arbor, z-stacks (1024×1024-pixel resolution, 1 µm per z-step) of neurons expressing fluorescent membrane markers were captured using a 20×0.7 NA (Leica SP5) 20×0.75 NA (Leica Stellaris) oil-immersion objectives. Maximum intensity projections of the z-stack images were created in FIJI, and neighboring neurons were cropped out using the freehand draw tool. These images were then further processed in FIJI by applying a threshold to eliminate background signal. The images were imported into Imaris, and Filament Tracer (BitPlane) with automatic detection was used to quantify total dendrite length and the number of terminal tips. The largest and smallest diameters of each neuron were manually measured to generate the dendrite start points and seed points. The thresholds were manually adjusted for the start points and seed points in order to cover the entire arbor and to reduce background points; seed points were manually added to segments that were not automatically identified. The filament was edited to remove the axon segment and to correct misdrawn segments. Measurements generated in Imaris were exported to Excel for further analysis.

### Quantification of FRAP

Analysis was performed in FIJI by creating maximum projections of the z-stacks from each time point. A line trace through the bleached region was drawn and the average intensity (arbitrary units; AU) of the center 10 µm was used to quantify the signal recovery over time. To account for general photobleaching, the signal intensity in the bleached 10 µm section was normalized by dividing by the average intensity (AU) of a 10 µm segment in a different secondary branch outside of the bleached region. Average intensity values were exported to Excel for further analysis. To calculate percent recovery of signal after photobleaching, the normalized average signal of the 10 µm branched region at 1, 3, 5, 10, and 20 minutes was divided by the initial signal from the pre-bleach z-stack.

### Statistical Analysis

All data were blinded prior to analysis. Statistical analysis was performed in Excel and GraphPad Prism using a significance level of p < 0.05. Outliers were identified using Grubbs’ test and removed. Data were analyzed for normality using the Shapiro-Wilk test. Normally distributed data were then analyzed for equal variance and significance using either an F-test and Student’s unpaired t-test (two samples) or one-way ANOVA with post-hoc Tukey (multiple samples). Data sets that were not normally distributed were analyzed using Mann-Whitney U test (two samples) or Kruskal-Wallis test with post-hoc Dunn test for significance (multiple samples). Significance levels are represented as follows: not significant (ns), p > 0.05; *, p = 0.05-0.01; **, p = 0.01-0.001; ***, p = 0.001-0.0001; and ****, p < 0.0001. Data are presented as the mean ± standard error of the mean (SEM) unless otherwise noted. In the graphs, *n* represents a neuron unless otherwise indicated.

## Acknowledgements

We thank Drs. Yuh Nung Jan (University of California, San Francisco), Hsiu-Hsiang Lee (National Taiwan University College of Medicine), Bing Ye (University of Michigan), the Bloomington Drosophila Stock Center (NIH P40OD018537), and the Vienna Drosophila Resource Center for fly strains and antibodies. We thank Dr. Anjon Audhya and Jennifer Peotter (University of Wisconsin-Madison) for assistance with the Imaris software. We thank members of the Wildonger lab for their feedback and suggestions on the project and manuscript; in particular, we thank Jessica Liang for her contributions to developing the FRAP protocol, Dena Johnson-Schlitz for technical assistance and advice, and Harriet Saunders for helpful guidance. We thank the Biochemistry Department and Dr. Aaron Hoskins (University of Wisconsin-Madison) for generously supporting J.M..

## Competing interests

The authors have declared that no competing interests exist.

## Author Contributions

Conceptualization: Josephine W. Mitchell and Jill Wildonger

Formal analysis: Josephine W. Mitchell and Ipek Midillioglu

Funding acquisition: Chun Han and Jill Wildonger

Investigation: Josephine W. Mitchell and Ipek Midillioglu

Resources: Bei Wang and Chun Han

Project administration: Chun Han and Jill Wildonger

Supervision: Jill Wildonger

Visualization: Josephine W. Mitchell

Writing – original draft: Josephine W. Mitchell and Jill Wildonger

Writing – review & editing: Josephine W. Mitchell, Ipek Midillioglu, Bei Wang, Chun Han, and Jill Wildonger

## Abbreviations

(DEG/ENaC/ASICs): Degenerin/Epithelial Na^+^ Channel/Acid Sensing Ion Channels
(da): dendritic arborization
(DN): dominant-negative
(dmn): dynamitin
(Dlic): Dynein light intermediate chain
(EcR): Ecdysone receptor
(ER): endoplasmic reticulum
(EC): extracellular
(FRAP): fluorescence recovery after photobleaching
(gRNA): guide RNA
(PI3K): phosphoinositide 3-kinase
(lva): lava lamp
(O/E): over-expressing
(Ppk): Pickpocket
(sfGFP): superfolder GFP
(secGFP): secreted GFP
(TRP): Transient Receptor Potential

## Supporting information

**S1 Fig. Effects of tagging endogenous Ppk1 on Ppk1 levels.**

**S2 Fig. Endogenous Ppk1 tagged with sfGFP at two different sites in an extracellular loop and Ppk1 tagged extracellularly with pHluorin.**

**S3 Fig. Characterization of a split-GFP approach to label membrane-expressed Ppk1.**

**S4 Fig. Characterization of GFP(11)EC::CD4::tdTomato.**

**S5 Fig. Aberrant Pickpocket channel activity disrupts dendrite morphogenesis.**

**S6 Fig. Disrupting Rab11 levels or activity causes a reduction in dendrite arbor growth.**

**S7 Fig. Wild-type Rab5 and Rab5-DN show a different pattern of distribution, and Rab5-DN reduces dendritic arbor morphogenesis.**

## Notes

### Competing Interest Statement

The authors have declared no competing interest.

### Summary of Updates

Addition of an author (Ethan Schauer) who made contributions to the work described in the manuscript.

