## Supplemental Figures for "Coordination of Pickpocket ion channel delivery and dendrite growth in Drosophila sensory neurons"

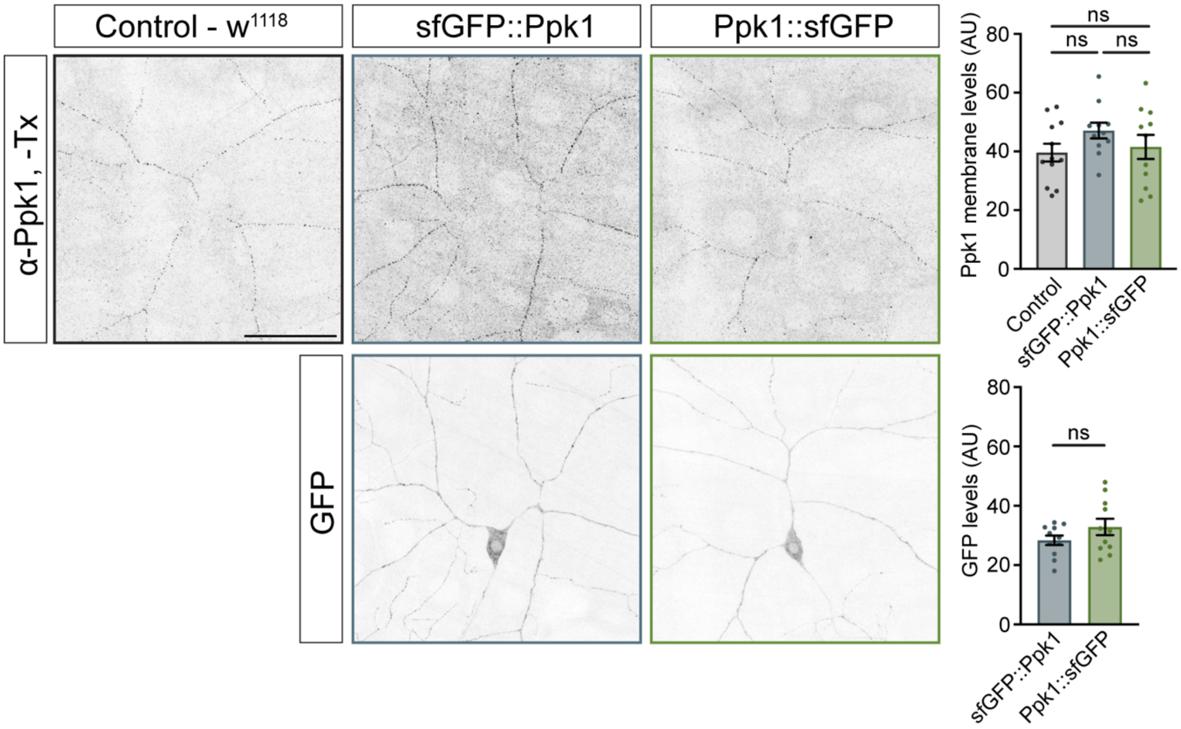


**S1 Fig. Effects of tagging endogenous Ppk1 on Ppk1 levels.** Representative images and quantification of membrane-expressed Ppk1, recognized by anti-Ppk1 antibodies (top) and sfGFP-tagged Ppk1 (bottom) in *w*$^{1118}$ control (12 larvae, 36 neurons), heterozygous N-terminally tagged Ppk1 (sfGFP::Ppk1) (11 larvae, 33 neurons), and heterozygous C-terminally tagged Ppk1 (Ppk1::sfGFP) (11 larvae, 33 neurons). Quantification, Ppk1 membrane levels: One-way ANOVA with post-hoc Tukey: *w*$^{1118}$ v. *sfGFP::Ppk1* (p=0.2570), *w*$^{1118}$ v. *Ppk1::sfGFP* (p=0.9097), *Ppk1::sfGFP* v. *sfGFP::Ppk1* (p=0.4802). Quantification, sfGFP-tagged Ppk1: Student's unpaired t-test (p=0.1707). In the graphs, each data point represents the average signal intensity per larva (2-3 neurons per larva). Data are plotted as mean ± SEM. n.s.=not significant (p>0.05). AU: arbitrary units. Scale bar, 50 µm.

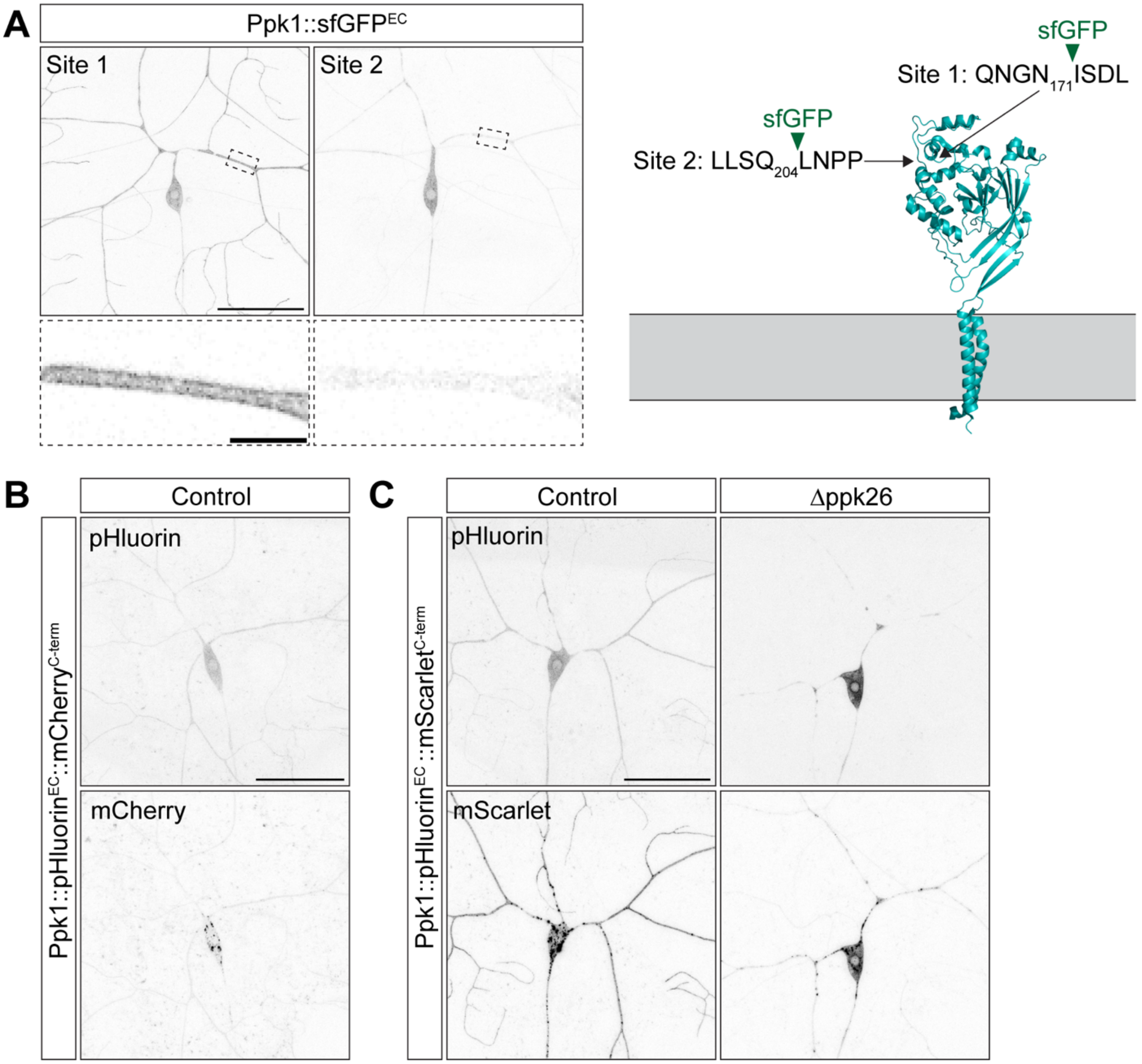


**S2 Fig. Endogenous Ppk1 tagged with sfGFP at two different sites in an extracellular loop and Ppk1 tagged extracellularly with pHluorin.** (A) Representative images of ddaC neurons expressing Ppk1 tagged with one copy of sfGFP at two different extracellular (EC) sites, Site 1 (between Asn171 and Ile172) and Site 2 (between Gln204 and Leu205). Ppk1 tagged with sfGFP at Site 1 showed similar fluorescent signal as Ppk1 tagged with sfGFP at the N- or C-terminus; thus, Site 1 was used for the insertion of additional tags [superecliptic pHluorin and GFP(11)]. Site 2 is located six amino acids downstream of the position at which a haemagglutinin (HA) tag was inserted (between F147 and K148) in rASIC1a (Chen and Gründer, 2007). On the right, a cartoon shows the crystal structure of an individual cASIC1 subunit; ASIC is comprised of three subunits (PDB: 2QTS; Jasti et al., 2007). The sites that correspond to where sfGFP was inserted in fly ppk1 are indicated (the fly amino acid sequences are shown). The locations of Site 1 and Site 2 were predicted by aligning the amino acid sequences of ppk1 and cASIC1. (B) Representative image of a ddaC neuron expressing Ppk1 tagged extracellularly (EC) with one copy of pHluorin at Site 1 and with mCherry at the C-terminus (Ppk1::pHluorin$^{EC}$::mCherry$^{C\text{-}term}$). The neuron is heterozygous for *Ppk1::pHluorin$^{EC}$::mCherry$^{C\text{-}term}$*. Scale bar, 50 µm. (C) Representative images of ddaC neurons expressing Ppk1 tagged extracellularly with one copy of pHluorin at Site 1 and with mScarlet at the C-terminus (Ppk1::pHluorin$^{EC}$::mScarlet$^{C\text{-}term}$) in *w$^{1118}$* control and *ppk26* null (Δppk26: *ppk26$^{\Delta 11/\Delta 11}$*) larvae. The neurons are homozygous for *Ppk1::pHluorin$^{EC}$::mScarlet$^{C\text{-}term}$*. Scale bar, 50 µm.

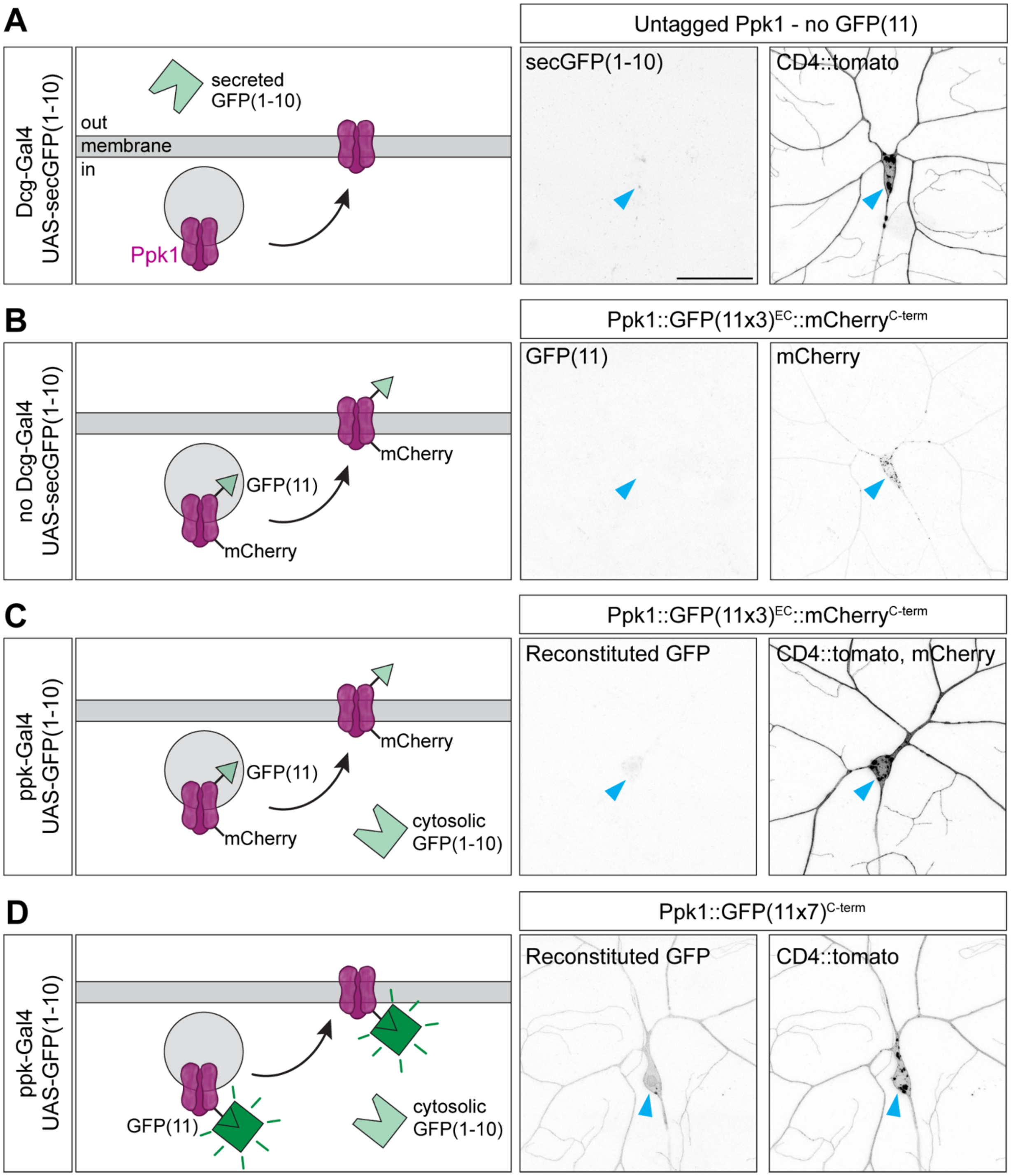


**S3 Fig. Characterization of a split-GFP approach to label membrane-expressed Ppk1.** (A) There is no GFP fluorescence in animals expressing *DcG-Gal4* and *UAS-secGFP(1-10)* in the absence of a protein tagged with GFP(11). (B) Without the *DcG-Gal4* driver, there is no GFP fluorescence in animals expressing *UAS-secGFP(1-10)* and *Ppk1::GFP(11x3)*$^{EC}$*::mCherry*$^{C\text{-}term}$. (C) There is no GFP fluorescence from Ppk1::GFP(11x3)$^{EC}$::mCherry$^{C\text{-}term}$ when GFP(1-10) is expressed in ddaC neurons by *ppk-Gal4*. The GFP(11x3) tag is positioned in an extracellular loop of Ppk1 and does not encounter cytosolic GFP(1-10). The red channel contains both the mCherry tag on the C-terminus of Ppk1 and a CD4::tdTomato membrane marker (*ppk-CD4::tdTomato*). (D) There is GFP signal from Ppk1 tagged at the C-terminus with GFP(11) [Ppk1::GFP(11x7)$^{C\text{-}term}$] when GFP(1-10) is expressed in ddaC neurons by *ppk-Gal4*. The GFP(11x7) tag is on the intracellular Ppk1 C-terminus, which enables GFP(11x7) to interact with cytosolic GFP(1-10). Scale bar, 50 μm.

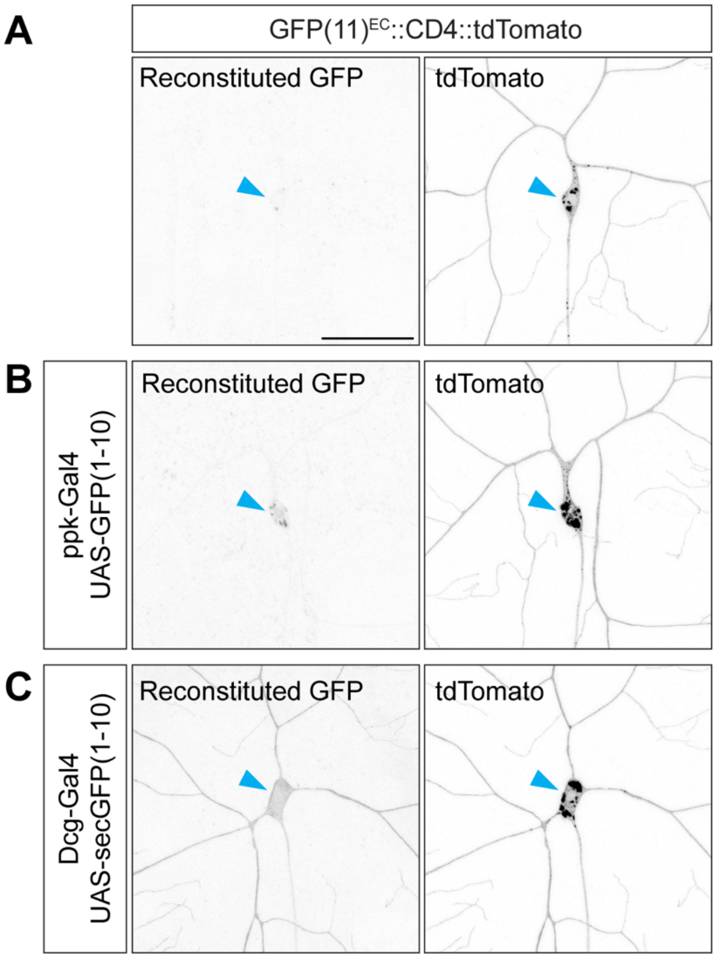


**S4 Fig. Characterization of conditions that allow for GFP fluorescence of GFP(11)$^{EC}$::CD4::tdTomato.** (A) There is no GFP fluorescence from GFP(11)$^{EC}$::CD4::tdTomato in the absence of secGFP(1-10). (B) There is no GFP fluorescence from GFP(11)$^{EC}$::CD4::tdTomato when GFP(1-10) is expressed in ddaC neurons by *ppk-Gal4*. The GFP(11) tag is positioned at the extracellular N-terminus of CD4 and does not encounter cytosolic GFP(1-10). (C) There is GFP signal from GFP(11)$^{EC}$::CD4::tdTomato when *DcG-Gal4* drives expression of *UAS-secGFP(1-10)*. A zoomed-in view of the cell body and proximal axon of this neuron is shown in Figure 2D. *GFP(11)$^{EC}$::CD4::tdTomato* is expressed in the class IV neurons under the control of the *ppk* enhancer (*ppk-GFP(11)$^{EC}$::CD4::tdTomato)*. Blue arrowheads point to the cell body. Scale bar, 50 μm.

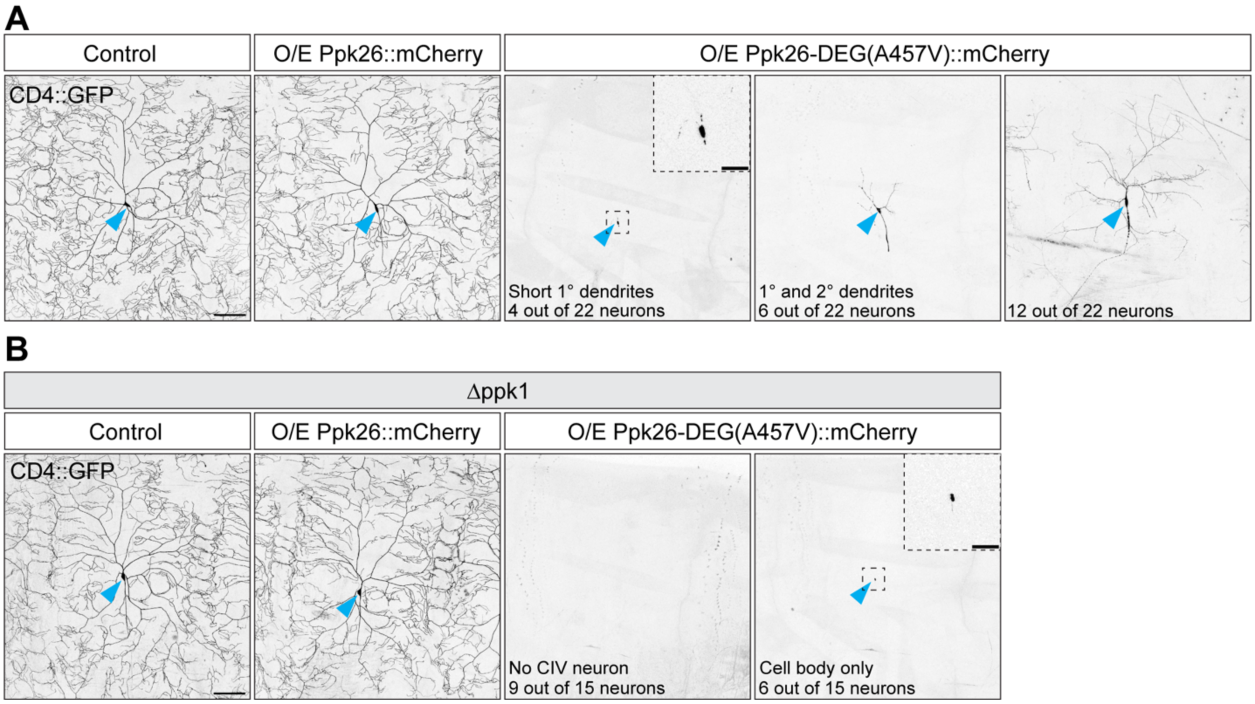


**S5 Fig. Aberrant Pickpocket channel activity disrupts dendrite morphogenesis.** Images of ddaC neurons in live 3rd instar larvae (120 h AEL). Neuron morphology was visualized with CD4::GFP (*ppk-CD4::GFP*). (A) Representative images of control neurons and neurons overexpressing wild-type Ppk26 (*UAS-Ppk26::mCherry*) or Ppk26 with the degenerin mutation [*UAS-Ppk26-DEG(A457V)::mCherry*]. *Ppk-Gal4* drove expression of the *UAS-Ppk26* constructs and was also included in the $w^{1118}$ control. The overexpression of Ppk26 with the degenerin mutation resulted in variable morphologies ranging from neurons with very short primary dendrites and no axon (left) to neurons with short primary and secondary dendrites (middle) to neurons with a severely reduced dendritic arbor (right). (B) Representative images of neurons in larvae lacking Ppk1 (Δppk1: $ppk1^{attP\text{-}KO/attP\text{-}KO}$) and, as indicated, overexpressing wild-type Ppk26 (*UAS-Ppk26::mCherry*) or Ppk26 with the degenerin mutation [*UAS-Ppk26-DEG(A457V)::mCherry*]. *Ppk-Gal4* drove expression of the *UAS-Ppk26* constructs and was also included in the Δppk1 control. In the absence of Ppk1, the overexpression of Ppk26 with the degenerin mutation resulted in the loss of ddaC neurons or ddaC neurons with a small cell body with no discernable axon or dendrites. Scale bars, 100 μm and 25 μm (dashed-outline boxes).

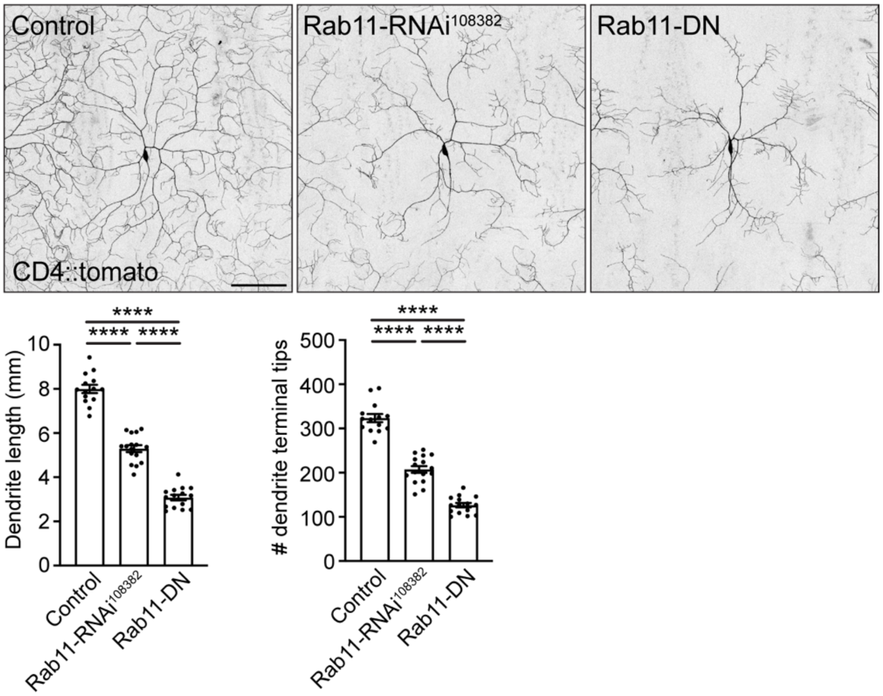


**S6 Fig. Disrupting Rab11 levels or activity causes a reduction in dendrite arbor growth.** Representative images of ddaC neurons in 72 h AEL larvae and quantification of dendrite length and dendrite tip number in control neurons ($w^{1118}$; 14 neurons, 6 larvae) and neurons over-expressing *Rab11-RNAi*$^{108382}$ (16 neurons, 7 larvae) or *Rab11-DN* (15 neurons, 7 larvae). Quantification, dendrite length: One-way ANOVA with post-hoc Tukey; $w^{1118}$ v. *Rab11-RNAi*$^{108382}$ ($p<0.0001$), $w^{1118}$ v. *Rab11-DN* ($p<0.0001$), *Rab11-RNAi*$^{108382}$ v. *Rab11-DN* ($p<0.0001$). Quantification, dendrite tips: One-way ANOVA with post-hoc Tukey: $w^{1118}$ v. *Rab11-RNAi*$^{108382}$ ($p<0.0001$), $w^{1118}$ v. *Rab11-DN* ($p<0.0001$), *Rab11-RNAi*$^{108382}$ v. *Rab11-DN* ($p<0.0001$). In the graphs, each data point represents a neuron, and data are plotted as mean ± SEM. The expression of *UAS-Rab11-RNAi*$^{108382}$ and *UAS-Rab11-DN* was driven by *ppk-Gal4*, and *ppk-Gal4* was included in the $w^{1118}$ control. Scale bar, 100 μm.

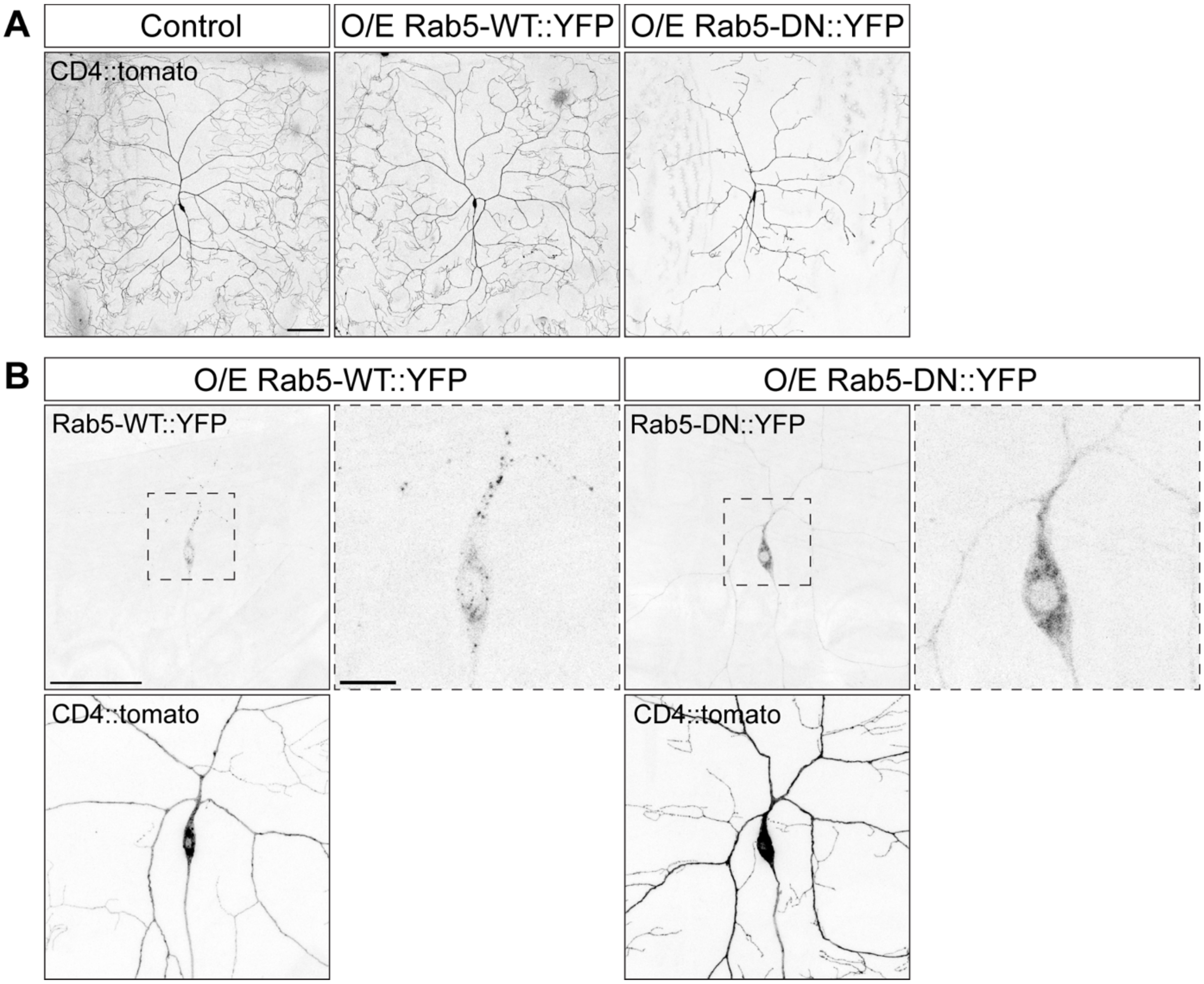


**S7 Fig. Rab5-WT and Rab5-DN show a different pattern of distribution, and Rab5-DN reduces dendritic arbor morphogenesis.** (A) Representative images of control ddaC neurons and ddaC neurons expressing Rab5-WT and Rab5-DN (*ppk-Gal4 UAS-Rab5-WT::YFP* and *ppk-Gal4 UAS-Rab5-DN::YFP*). The neuronal membrane is marked by CD4::tdTomato (*ppk-CD4::tdTomato*). Scale bar, 100 μm. (B) Representative images showing the distribution of Rab5-WT::YFP and Rab5-DN::YFP in ddaC neurons from fixed larval fillets with zoomed-in images of the cell bodies (dashed-outline boxes). The neuronal membrane is marked by CD4::tdTomato (ppk-CD4::tdTomato). Scale bar, 50 μm and 10 μm (dashed-outline box).
